# Auditory processing remains sensitive to environmental experience during adolescence

**DOI:** 10.1101/2021.04.12.439537

**Authors:** Kelsey L. Anbuhl, Justin D. Yao, Robert A. Hotz, Todd M. Mowery, Dan H. Sanes

**Author notes:** Contact: Kelsey L. Anbuhl; Dan H. Sanes. **Author Contributions** KLA and DHS designed research and secured funding; KLA collected behavioral and *in vivo* physiological data; RAH collected behavioral control data in Figure 9; TMM collected *in vitro* physiology data; KLA, TMM, and JDY analyzed data. KLA and DHS wrote the manuscript.

## Abstract

Development is a time of great opportunity. A heightened period of neural plasticity contributes to dramatic improvements in perceptual, motor, and cognitive skills. However, developmental plasticity poses a risk: greater malleability of neural circuits exposes them to environmental factors that may impede behavioral maturation. While these risks are well-established prior to sexual maturity (i.e., critical periods), the degree of neural vulnerability during adolescence remains uncertain. To address this question, we induced a transient period of hearing loss (HL) spanning adolescence in the gerbil, confirmed by assessment of circulating sex hormones, and asked whether behavioral and neural deficits are diminished. Wireless recordings were obtained from auditory cortex neurons during perceptual task performance, and within-session behavioral and neural sensitivity were compared. We found that a transient period of adolescent HL caused a significant perceptual deficit (i.e., amplitude modulation detection thresholds) that could be attributed to degraded auditory cortex processing, as confirmed with both single neuron and population-level analyses. In contrast, perceptual deficits did not occur when HL of the same duration was induced in adulthood. To determine whether degraded auditory cortex encoding was attributable to an intrinsic change, we obtained auditory cortex brain slices from adolescent HL animals, and recorded synaptic and discharge properties from auditory cortex pyramidal neurons. There was a clear and novel phenotype, distinct from critical period HL: excitatory postsynaptic potential amplitudes were elevated in adolescent HL animals, whereas inhibitory postsynaptic potentials were unchanged. This is in contrast to critical period deprivation, where there are large changes to synaptic inhibition. Taken together, these results show that diminished adolescent sensory experience can cause long-lasting behavioral deficits that originate, in part, from a dysfunctional cortical circuit.

Summary of experimental design and main findings.

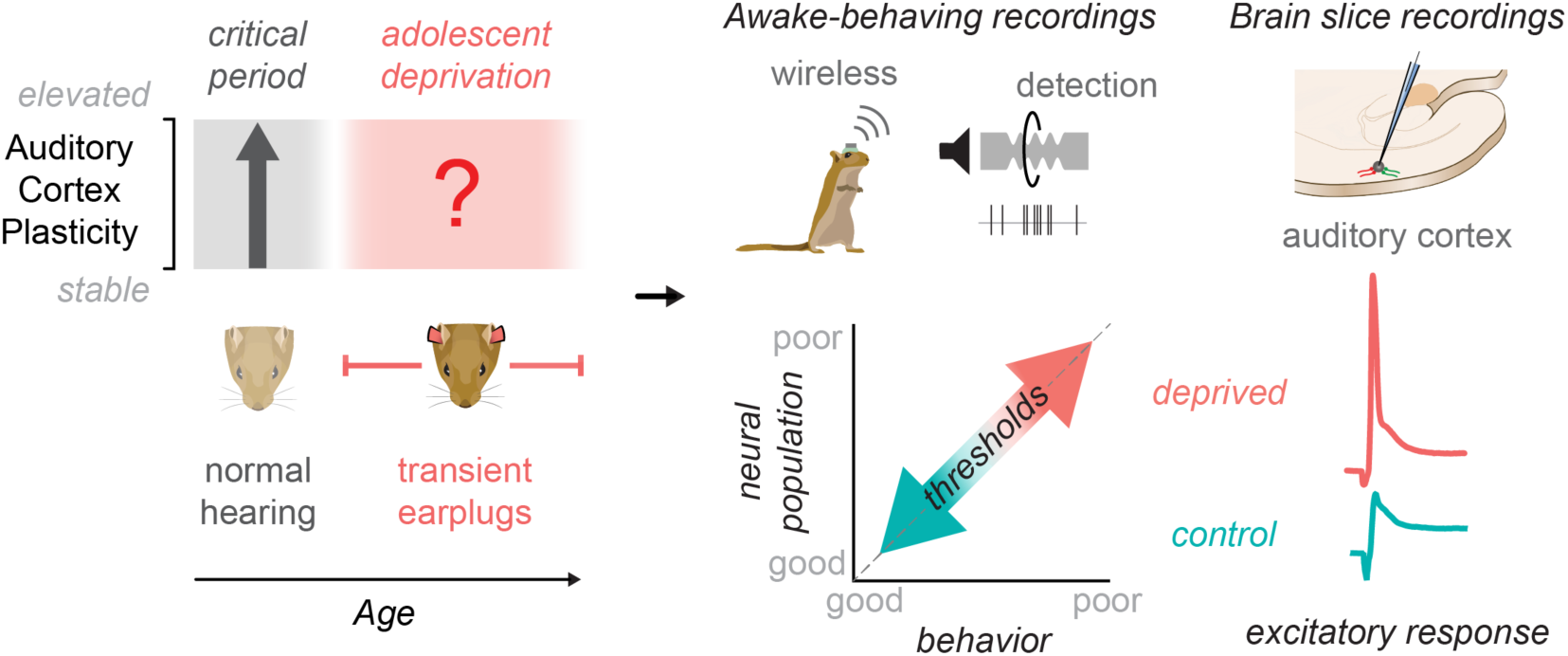

## Main text / Introduction

The adolescent brain is distinguished by structural and functional changes that coincide with late-developing behavioral skills. In humans, synaptic pruning, myelination and cortical evoked-potentials each continue to mature well into the second or third decade of life (Sharma et al., 1997; Giedd et al., 1999; Moore and Guan, 2001; Bishop et al., 2007; Sussman et al., 2008; Pinto et al., 2010; Lebel and Beaulieu, 2011; Petanjek et al., 2011; Mahajan and McArthur, 2012b, 2012a; Shafer et al., 2015). Similarly, many behavioral skills are late to mature, including auditory perceptual learning (Huyck & Wright, 2011, 2013), temporal processing skills required for aural language (Banai et al., 2011; McMurray et al., 2018), face recognition (Carey et al., 1980; Germine et al., 2011), and fine motor control (Dayanidhi et al., 2013). Furthermore, the transition from childhood to adulthood is characterized by prolonged development of executive function (Selemon, 2013; Downes et al., 2017; Simmonds et al., 2017), emotion regulation (Guyer et al., 2008; Cohen Kadosh et al., 2013; Kadosh et al., 2013), and social skills (Blakemore, 2012). At the same time, adolescence is associated with risk factors, such as stress (Eiland and Romeo, 2013) or substance abuse (Davidson et al., 2015), and is a time during which many psychiatric disorders emerge (Paus et al., 2008). Here, we ask whether adolescent plasticity poses a risk for neural and behavioral development of sensory function, similar to postnatal critical periods (Hubel and Wiesel, 1970; Van der Loos and Woolsey, 1973; Knudsen et al., 1984b; Hensch, 2005; de Villers-Sidani et al., 2007; Popescu and Polley, 2010; Cheetham and Belluscio, 2014; Mowery et al., 2015).

There is broad agreement that, prior to sexual maturity, sensory deprivation can permanently disrupt central nervous system function when initiated during brief epochs, termed developmental critical periods (Hubel and Wiesel, 1970; Van der Loos and Woolsey, 1973; Knudsen et al., 1984b; Hensch, 2005; de Villers-Sidani et al., 2007; Popescu and Polley, 2010; Cheetham and Belluscio, 2014; Mowery et al., 2015). For example, a brief period of mild hearing loss (HL) can induce long-lasting changes to gerbil auditory cortex inhibitory synapses when it occurs before postnatal day (P) 19, but not after (Mowery et al., 2015, 2017). Furthermore, a similar period of HL induces perceptual deficits (Caras and Sanes, 2015), which can be rescued by restoring synaptic inhibition (Mowery et al., 2019). A perceptual deficit has also been reported for 5-year-old children with a history of transient HL (McKenna Benoit et al., 2018), in agreement with other human studies suggesting critical periods prior to sexual maturity (Sharma et al., 2002; Svirsky et al., 2004; Putzar et al., 2007, 2010).

While non-human research has focused on early critical periods, childhood HL can often emerge after birth (Lü et al., 2011; Barreira-Nielsen et al., 2016) and extend through adolescence (Niskar et al., 2001; Shargorodsky et al., 2010), resulting in more severe language deficits than those with brief HL (Yoshinaga-Itano et al., 1998; Tomblin et al., 2015). In fact, a majority of adolescents exhibit a mild-minimal high-frequency hearing loss that coincides with poorer speech perception in noise (Zadeh et al., 2019). Adolescents with HL are at risk for social isolation (Patel et al., 2020), and higher rates of psychiatric, depressive, or anxiety disorders (Theunissen et al., 2014). Thus, adolescence may be associated with greater vulnerability to even a transient period of auditory deprivation. To address this question, we asked whether a transient period of auditory deprivation, beginning after the auditory cortex (AC) critical period closes (Mowery et al., 2015, 2017, 2019) and extending throughout the period of sexual maturation, disrupts perceptual and neural auditory function. By comparing this to an identical manipulation that began in adulthood, we found that a temporary period of adolescent HL selectively impaired detection of amplitude modulations (AM), a foundational sound cue for aural communication including speech (Singh and Theunissen, 2003). These perceptual deficits were linked to poorer AC neuron encoding during task performance, and a change to AC synaptic properties distinct from that observed following critical period deprivation (Mowery et al., 2015, 2017, 2019). Taken together, our study suggests a sensitive period for sensory function during adolescent development.

## Results

### Transient sensory deprivation spans adolescence

To determine whether auditory function is vulnerable during adolescence, we induced transient hearing loss (HL) at postnatal (P) day 23, after a well-defined AC critical period ends (Mowery et al., 2015). Normal hearing was restored (i.e., earplugs were removed) at P102, after the animals had passed through adolescence and reached sexual maturity (Figure 1A). To confirm the time course of sexual maturation, we tracked testosterone levels across development, both in normal hearing animals (n=12) and littermates with transient auditory deprivation during adolescence (n=14). Figure 1B shows serum testosterone levels for male (solid line) and female (dotted line) gerbils from P35 to P102. For males, testosterone levels were negligible at P35 (<0.2 ng/mL), rose sharply at P54 (2.42±0.8 ng/mL), and remained elevated thereafter. A two-way repeated measures ANOVA revealed a significant effect of age on testosterone levels (*F*_*(4,39)*_ = 3.7, p=0.01), no effect of hearing loss (*F*_*(1,39)*_ = 0.07, p=0.8), and no interaction between the two variables (*F*_*(4,39)*_= 0.56, p=0.7). For females, there was a small, transient rise in testosterone between P35 and P54, with no significant effect of hearing loss on testosterone levels (*F*_*(1,38)*_ = 3.13, p=0.08). Therefore, the auditory deprivation between P23-102 likely spans the entirety of adolescence. Furthermore, we examined estradiol, another sex hormone, in normal hearing and HL animals at P90-102. We found comparable levels of serum estradiol in the females of both groups (Ctrl: 0.004 ng/mL; HL: 0.005 ng/mL) and negligible amounts in the males of both groups (Ctrl: <0.003 ng/mL; HL: <0.003 ng/mL). This suggests that both normal hearing and HL groups had similar levels of sex hormones by P102, the age at which the earplugs were removed (for the HL group).

**Figure 1.**
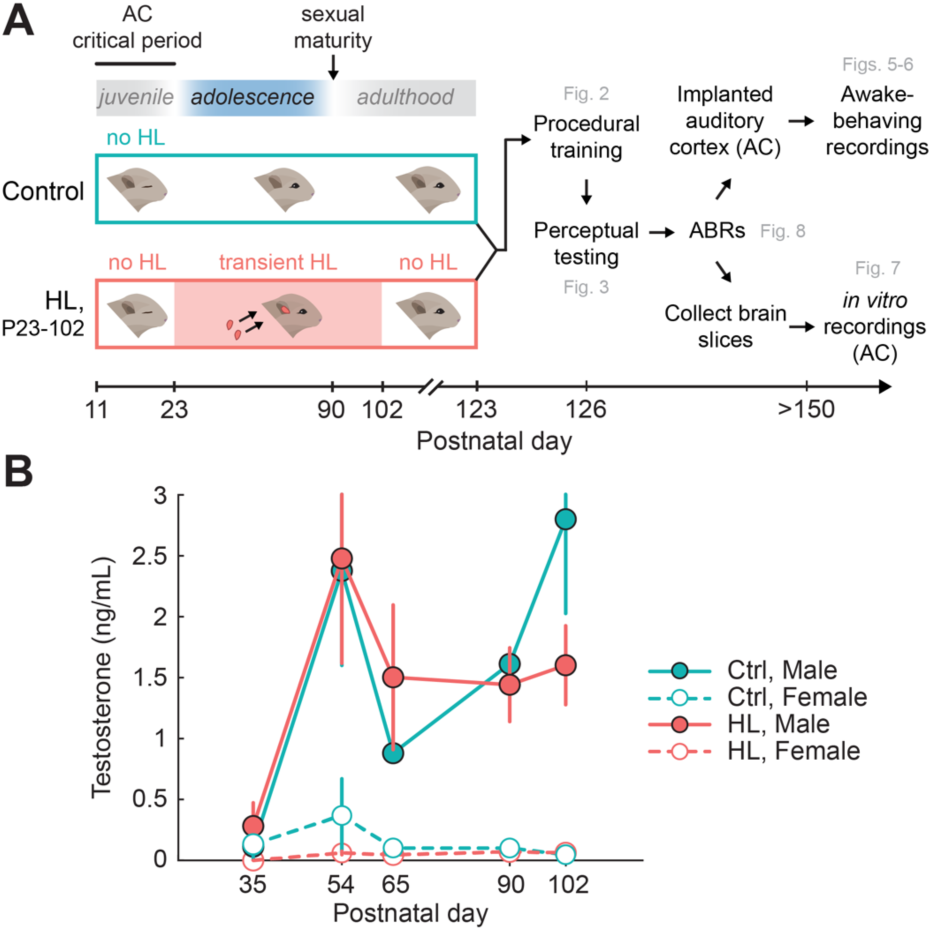
Transient sensory deprivation during adolescence. **(A)** Experimental timeline of manipulation and behavioral, physiological assessment. Gerbils either received transient hearing loss (HL; via bilateral earplugs) from postnatal (P) day 23, after the auditory cortex (AC) critical period ends, through P102, after sexual maturity (n=14) or no earplugs (littermate controls, n=12). Following earplug removal, animals recovered for 21 days prior to behavioral training and perceptual testing. Following behavioral assessment, auditory brainstem responses (ABRs) were collected in all animals. After ABR collection, animals were either chronically implanted in the AC and used for awake-behaving recordings (Ctrl: n=7; HL: n=9), or had brain slices collected for *in vitro* AC recordings (Ctrl: n=5; HL: n=5). **(B)** Serum testosterone levels across age for male (solid line) and female (dotted line) gerbils. The auditory deprivation induced spanned the entire time course of sexual maturation (estradiol not shown).

### Adolescent hearing loss did not alter procedural learning

Following transient HL and earplug removal at P102, gerbils recovered for 21 days prior to behavioral training on the amplitude modulation (AM) depth detection task (Figure 2A). Control and adolescent HL animals learned the AM detection task procedure at a similar rate (Figure 2B). A two-way mixed-model ANOVA revealed a significant effect of trial number on performance (F_(74,1776)_ = 19.8, p<0.0001), but no effect of earplug experience (F_(1,24)_ = 0.2685, p = 0.61), and no interaction between the two variables (F_(74, 1776)_ = 1.18, p = 0.1413). Thus, normal hearing controls and adolescent HL animals required a comparable number of Warn trials to reach the criterion for training performance (d’ ≥ 1.5; Figure 2C). A one-way ANOVA revealed that the HL manipulation had no effect on the number of trials required to reach criterion during training (*t*_*(24)*_ = −0.55; p = 0.59). In addition, the HL manipulation had no effect on the average d’ achieved during the last 20 trials of procedural training (one-way ANOVA: *t*_*(24)*_ = −0.23; p = 0.82; Figure 2D). Therefore, the two groups were equally proficient at task performance prior to psychometric testing.

**Figure 2.**
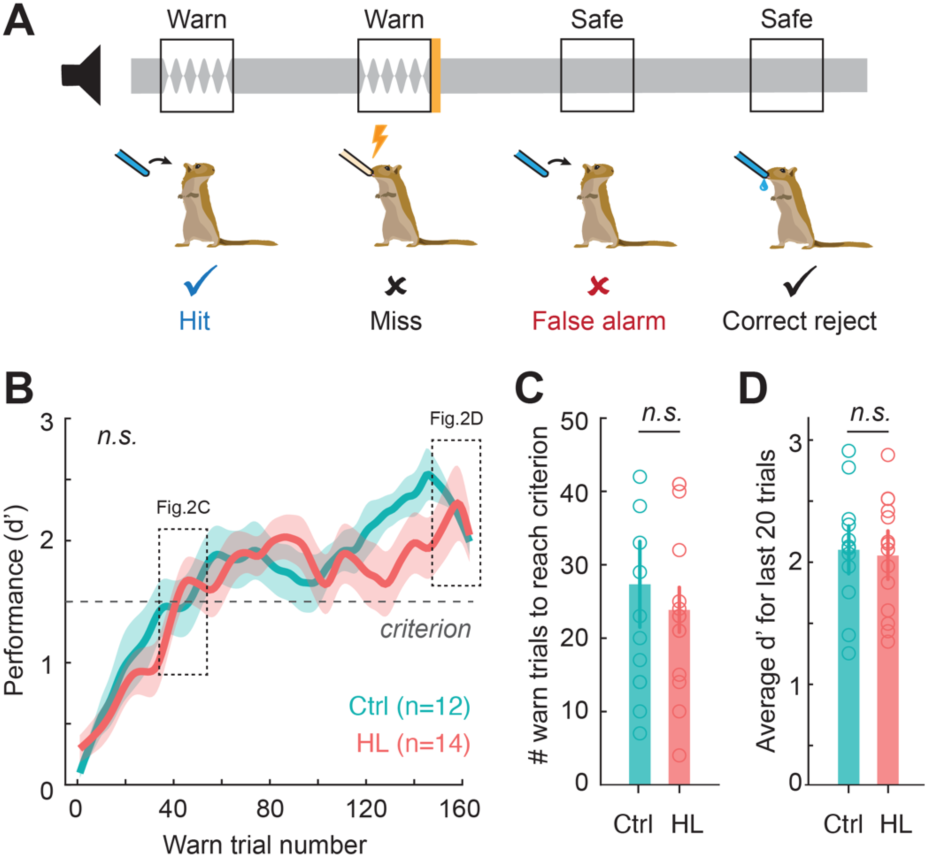
Hearing loss (HL) during adolescence did not alter procedural training performance in adulthood. **(A)** Schematic of the amplitude modulation (AM) detection task. Animals were trained to drink from a water spout in the presence of continuous broadband noise (Safe trials; classified as a “correct reject”) and to cease drinking when the noise transitioned to a modulated “Warn” signal (0 dB relative to 100% AM noise; 5 Hz rate; classified as a “Hit”). Failure to withdraw from the spout during Warn trials (classified as a ‘‘Miss”) resulted in a mild aversive shock. Withdrawing from the spout during Safe trials is classified as a “False Alarm”. Behavioral performance is quantified by utilizing the signal detection metric, d’. **(B)** Procedural training sessions (3-4 separate sessions) were combined to compute behavioral performance (d’) as a function of warn trial number using a 5-trial sliding window. Dotted line boxes correspond to data shown in C, D. Data are depicted as the mean ± SEM. **(C)** Number of warn trials to reach performance criterion (d’≥1.5; see horizontal line in **B)** for control (Ctrl) and HL-reared animals. **(D)** Average d’ for the last 20 trials of procedural training for each group.

### Adolescent hearing loss impaired amplitude modulation detection thresholds

AM depth detection thresholds were assessed over 10 separate days of psychometric testing. Figure 3A shows psychometric functions on the first day of perceptual testing for normal-hearing animals (n=12) and those that experienced adolescent HL (n=14). HL animals displayed significantly poorer AM detection thresholds on the first day of testing (Ctrl: −11±0.5, HL: −8±0.7 dB re: 100% AM; *t*_(24)_ = 2.98, *p* = 0.007; Figure 3B). The AM detection deficit did not resolve with perceptual learning (Sarro and Sanes, 2010, 2011; Fitzgerald and Wright, 2011; Caras and Sanes, 2015, 2017, 2019) over 10 consecutive sessions. In fact, adolescent HL animals displayed poorer AM detection thresholds across all testing days, even though both groups exhibited perceptual learning (*initial*: HL: −8 dB; Ctrl: −11 dB re: 100% AM; *final*: HL: −13 dB; Ctrl: −16 dB re: 100% AM; Figure 3B). An analysis of covariance (ANCOVA; hearing status × log[test day]) revealed a significant effect of task experience (*F*_*(1,206)*_ = 58.1, p<0.0001), such that AM detection thresholds decreased by 5 dB (re: 100% AM depth) per log(test day). There was no significant interaction between log(test day) and hearing status (*F*_*(1,206)*_ = 0.72, p=0.4), but there was a significant effect of HL alone (*F*_*(1,206)*_ = 53.2, p<0.0001) with AM detection thresholds 2.9 dB higher than controls across all testing days.

**Figure 3.**
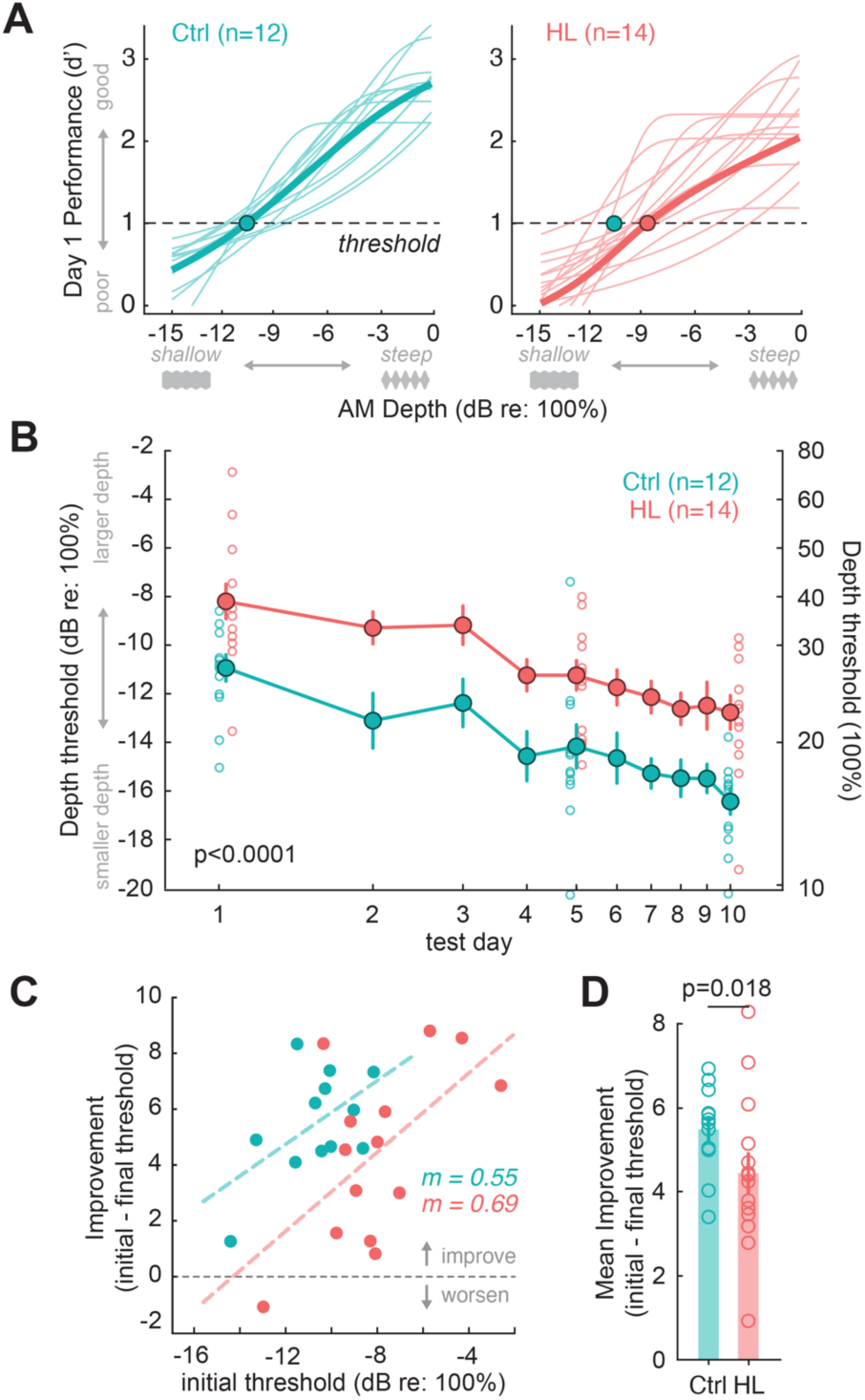
Transient hearing loss during adolescence impairs amplitude modulation (AM) depth detection in adulthood. **(A)** Psychometric functions on the first day of perceptual testing for control (Ctrl, n=12) and hearing loss (HL, n=14) animals. Threshold was defined as the depth at which d’=1 (dotted line). AM depths are presented on a dB scale (re: 100% depth), where 0 dB corresponds to 100% modulation, and decreasing values indicate smaller depths (e.g., see grey depth stimuli visualized below x-axes. Circle indicates the average group detection threshold. **(B)** HL animals display significantly poorer AM detection thresholds than controls on the first day of testing (Ctrl: −11 ±0.5, HL: −8±0.7 dB re: 100%; p=0.007). The AM detection deficit for HL animals persisted across 10 consecutive testing days. An analysis of covariance (ANCOVA) revealed the effect of HL alone to be significant (p<0.0001). Filled circles and lines indicate the group mean±SEM. Open circles indicate individual data points for the first, middle, and last psychometric testing session to highlight between-subject variablity between control and adolescent HL animals. **(C)** Perceptual improvement (initial subtracted from final threshold) as a function of initial threshold. Values above the dotted line indicate an improvement in thresholds, whereas values below indicate a worsening of thresholds. Dotted lines indicate fitted linear regressions (R^2^=0.3-34). The slopes (m) of the regression fits were not significantly different from one another (p=0.8), whereas the y-axis intercepts were significantly different indicating the average amount of improvement was lower in adolescent-HL animals (p<0.0001). **(D)** The mean improvement from the initial to the final session for each group. Control animals improved significantly more than transient HL animals (ANCOVA; p=0.018).

Elevated detection thresholds could be attributed to poorer hit rates (i.e., fewer correct responses to AM Warn trials) and/or elevated false alarm (FA) rates (i.e., incorrectly withdrawing from spout during Safe trials). Here, we find that FA rates consistently remain very low (< 0.06) across all testing sessions for both control and HL animals (grand average across testing sessions, Ctrl: 0.05; HL: 0.06; FA data not shown). An ANCOVA reveals no significant effect of testing day on FA rate (*F*_*(1,254)*_ = 0.01, p=0.92), and no effect of HL (*F*_*(1,254)*_ = 1.15, p=0.28). Therefore, the elevated detection thresholds in adolescent HL animals are due to poorer hit rates during AM Warn trials (Hit rate data not shown).

To determine whether there was a significant difference in perceptual learning, we plotted perceptual improvement across the 10 testing days (i.e., initial threshold – final threshold) as a function of each animal’s initial detection thresholds (Figure 3C). The slopes of the linear regression fits to each group were not significantly different from one another (ANCOVA: *F*_*(1,21)*_ = 0.07, p=0.8), but the adolescent HL curve had a significantly smaller y-axis intercept parameter (p<0.0001), showing that adolescent HL impaired perceptual learning. Figure 3D compares the mean improvement over 10 days of practice: Controls improved significantly more than transient HL animals (ANCOVA; initial threshold as a covariate; df=1, F=6.5769, p=0.018).

Given the role that sex hormones play in the development and maintenance of auditory pathways (e.g., Canlon and Frisina, 2009), and the between-sex differences in hormones measured in this study (Figure 1), we examined whether there were sex-specific differences in procedural learning or psychometric performance that might account for some of the between-subject variability observed in Figures 2-3. Therefore, we plotted the procedural training data separately for the males and females within each group, we find no significant differences in the rate at which males or females learned the task procedure (Supplemental Figure 1A; mixed-model ANOVA: *F*_*(1,4)*_ = 0.144, *p* = 0.72). Further, males and females within each group required a comparable number of Warn trials to reach criterion for training performance (d’ ≥ 1.5; Supplemental Figure 1B; mixed-model ANOVA: *F*_*(3, 14)*_ = 0.212, *p* = 0.89), and achieved comparable performance (average d’) during the last 20 trials of procedural training (Supplemental Figure 1C; mixed-model ANOVA: *F*_*(3, 14)*_ = 0.33, *p* = 0.14). Therefore, we did not find sex-specific differences in procedural training on the behavioral task. Next, we examined whether there were male and female differences in perceptual thresholds across the 10 days of psychometric testing. A two-way mixed-model ANOVA followed by a Tukey’s test for multiple comparisons revealed no significant effect of sex in control animals (*p* = 0.43) or in HL animals (*p* = 0.97). See Supplemental Figure 1D for depth detection thresholds on the final day of testing for male and female Control and HL animals.

Chronic stress during adolescence can have adverse effects on central nervous system structure and function (Isgor et al., 2004). To determine whether the perceptual deficits that we observed were due, in part, to elevated stress associated with earplugs, we obtained cortisol levels in each animal (Supplemental Figure 2) at five timepoints across development (P35, P55, P65, P90, and P102). We found no significant differences in cortisol levels between normal-hearing and adolescent HL animals at any of the ages tested (Supplemental Figure 2A). In addition, we found no significant correlations between cortisol level at any developmental age with perceptual performance on the first day of testing (Pearson’s *r* = −0.4 – 0.3; p-values of linear regression = −0.2-0.7), validating that the deficits observed were not due to elevated early life stress (Supplemental Figure 2B).

### Adult-onset hearing loss did not alter procedural learning or amplitude modulation detection

Next, we asked whether the perceptual deficits observed following adolescent hearing loss was specifically due to the developmental period, or whether the long duration of deprivation itself caused the effect. To test this, we induced the same duration of HL (80 days) beginning in adulthood (≥P102) in a separate set of animals (Adult controls, n=8; Adult-onset HL, n=8). Following 80 days of HL, earplugs were removed and animals experienced the same duration of recovery (3 weeks) prior to training and testing on the AM depth detection task. Adult controls and adult-onset HL animals learned the task procedure at the same rate (Figure 4A). A two-way mixed model ANOVA revealed a significant effect of trial number on performance (*F*_*(159,318)*_ = 3.18, p<0.001), no effect of earplug experience (*F*_*(1,2)*_ = 0.002, p=0.97), and no interaction between the two variables (*F*_*(159, 318)*_ = 0.96, p=0.62). Both groups took the same number of warn trials to reach criterion for training performance (Figure 4B; *F*_*(1,14)*_ = 0.0013 p = 0.97), and both groups achieved comparable performance (d’) on the last 20 trials of procedural training prior to psychometric testing (Figure 4C; *F*_*(1,14)*_ = 0.61 p = 0.45).

**Figure 4.**
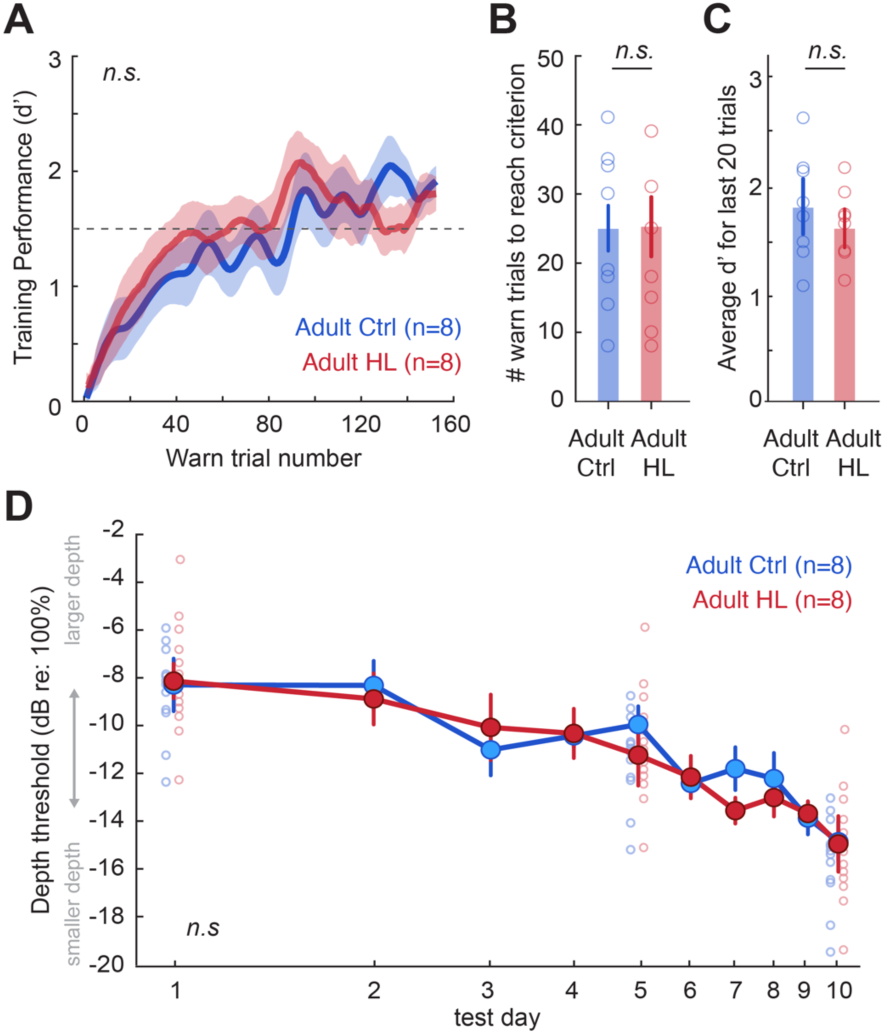
Adult-onset hearing loss (>P102) of the same duration (80 days) does not impair procedural training or amplitude modulation (AM) depth detection following earplug removal and recovery. **(A)** Procedural training sessions (3-4 separate sessions) were combined and behavioral performance (d’) was computed as a function of warn trial number using a 5-trial sliding window. Data for Adult controls (blue; n=8) and Adult-onset HL (red; n=8) are depicted as the mean ± SEM. **(B)** Number of warn trials to reach performance criterion (d’≥1.5). **(C)** Average d’ for the last 20 trials of procedural training prior to psychometric testing. **(D)** Psychometric functions were collected for 10 consecutive testing days for Adult controls and Adult-onset HL animals. Detection thresholds, defined as the depth at which d’=1, were computed for each testing session. AM depths are presented on a dB scale (re: 100% depth), where 0 dB corresponds to 100% modulation, and decreasing values indicate smaller depths. Circles indicate the average group detection threshold (±SEM) and lines show individual animal performance. Both groups exhibit comparable thresholds on the first day of testing (Adult Ctrl: −8.33±1; Adult HL: −8.17±0.7 dB re: 100%) and last day of testing (Adult control: −14.83±0.8; Adult-onset HL: −14.9±1.1 dB re: 100% AM). An analysis of covariance (ANCOVA) revealed a significant effect of testing day (p<0.0001), but no significant effect of Adult-onset HL alone (p=0.87).

AM depth detection thresholds were then assessed over 10 consecutive days of psychometric testing for Adult controls and Adult-onset HL animals (Figure 4D). Both groups exhibited nearly identical detection thresholds on the first day of testing (Adult control: −8.33±1.09; Adult-onset HL: −8.17±0.7 dB re: 100% AM) and the final testing day (Adult control: −14.83±0.8; Adult-onset HL: − 14.9±1.1 dB re: 100% AM). An analysis of covariance (ANCOVA; hearing status × log[test day]) revealed a significant effect of testing day (*F*_*(1,154)*_ = 84.3, p<0.0001), but no significant interaction between log(test day) and hearing status (*F*_*(1,154)*_ = 0.25, p=0.62), and no significant effect of HL alone (*F*_*(1,154)*_ = 0.03, p=0.87). This in stark contrast to the detection and perceptual learning deficits that were observed following the same HL duration in adolescence. Therefore, we attribute the perceptual deficits observed to adolescent vulnerability, and not the long duration of deprivation itself.

### Diminished auditory cortex (AC) neuron detection thresholds during task performance in adolescent HL animals

Since perceptual deficits persisted long after peripheral input was restored, we asked whether it could be attributed to impaired auditory cortex (AC) encoding. A subset of behaviorally-tested gerbils (Ctrl: n=7; HL: n=9) were implanted with chronic 64 channel silicone electrode arrays in the AC, and wireless neural recordings were collected as they performed the AM detection task (Figure 5A-B). We recorded from a total of 2,183 multi- and single-cortical units (Ctrl: n=846; HL: n=1337). Since individual sensory neurons have been shown to be predictive of behavior in perceptual discrimination (Pitkow et al., 2015) and detection tasks (Caras and Sanes, 2017), we opted to first examine detection thresholds for individual auditory cortical neurons. Multi- and single-units were selected for further analysis if they met the criteria for AM responsiveness (see Methods). We found 265 AM responsive units from control animals (multi-units: n=188; single-units: n=77) and 216 AM responsive units from adolescent HL animals (multi-units: n=154; single-units: n=62). The relatively low yield of AM responsive units included in the analysis allowed us to determine whether sensory encoding of AM can be explained by the sparse coding of a small subset of neurons and whether those neurons drive perceptual ability. Figure 5C shows raster plots and corresponding post-stimulus time histograms (PSTHs) for two example single units collected from a control and an adolescent HL animal. The plots display neural responses to unmodulated noise (Safe signal; *bottom panel*) and to a range of AM depths (Warn signal), from steeply modulated (−6 dB re: 100%; *top panel*) to shallow modulations (−18 dB re: 100%). Both example neurons display an increase in firing rate activity for modulated trials compared to unmodulated trials. In addition, firing rate increases with larger AM depths (Supplemental Figure 3A).

**Figure 5.**
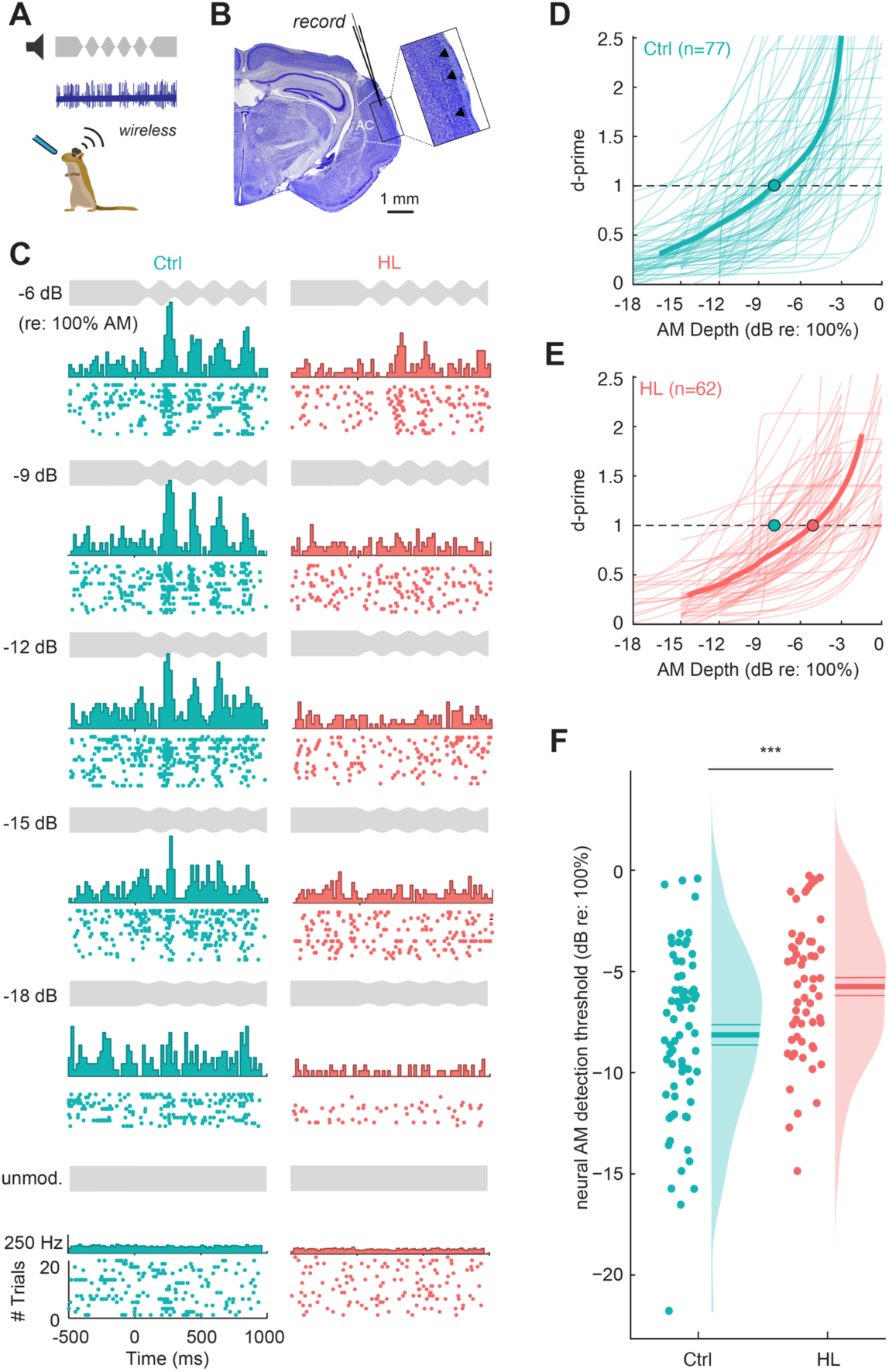
Single-unit analysis reveals poorer neural detection thresholds in the auditory cortex. **(A)** Chronic 64 channel electrode arrays were implanted into the auditory cortex (AC) of a subset of control (n=7) and HL (n=9) animals, and wireless neural recordings were collected as they performed the AM depth detection task. **(B)** Representative Nissl-stained coronal section from one implanted animal. Inset shows electrode track through primary AC (see arrows). **(C)** Raster plots with poststimulus time histograms are shown for two example single units from a control (left panel; Subject ID F276484) and HL (right panel; Subject ID M277481) animal. Plots are arranged in order from larger AM depths (top) to smaller depths (towards bottom; see depth stimulus in grey for reference). **(D, E)** The firing rate of single units that met the criteria for AM sensitivity were transformed into d’ values (see Methods). Neural d’ values were fit with a logistic function and plotted as a function of AM depth for a population of control (n=77) and HL (n=62) single units. Thin lines indicate individual fits, and the thick line indicates the population mean of the fits. Neural thresholds were defined as the AM depth at which the fit crossed d’=1. Circle indicates the mean neural depth threshold for each population. **(F)** Neural thresholds are plotted for control and HL animals. Individual thresholds are shown (circles), along with a half-violin plot indicating the probability density function. Horizontal lines indicate the mean±SEM. Single units from HL animals exhibit poorer neural depth thresholds than control single units (p = 0.0007).

To determine whether adolescent HL alters AM detection thresholds, the firing rate of multi- and single-units that met the criteria for AM responsivity were transformed into d’ values (see Methods). Neural d’ was plotted as a function of AM depth and fit with a logistic function to generate neurometric functions for each cell. Figure 5D,E shows neurometric functions for a population of single units from control and HL animals. Similar to psychometric functions, neural detection thresholds were defined as the depth at which the neurometric fit crossed a *d’=1*. AC neurons from HL animals displayed significantly poorer thresholds, as compared to control neurons (Ctrl: −8.01±0.5; HL: −5.65±0.5 dB re: 100%; Figure 5F). A one-way ANOVA reveals a significant effect of HL on single unit neural thresholds (*F*_*(1,128)*_ = 12.1, *p* = 0.0007). We find a similar effect of adolescent HL when we examine multi-unit neural thresholds (Ctrl: −6.8±0.2; HL: −5.7±0.2 dB re: 100%; *F*_*(1,321)*_ = 8.71, *p* = 0.003; Supplemental Figure 3F). A two-way mixed model ANOVA reveals a significant effect of hearing status on neural thresholds (*F*_*(1,449)*_ = 23.61, *p*<0.001), but no effect between unit type (single- or multi-units; *F*_*(1,449)*_ = 2.85, *p*=0.09). Though only the “best” neurons were included in the analysis (i.e., met strict criteria for AM-responsivity), the AC neurons from adolescent HL animals still display poorer neural detection to AM.

Poorer AC neuron detection thresholds could be attributed to alterations in basic response properties to AM depth stimuli. We found no effect of HL on the firing rate of single-units to the depths presented (two-way mixed-model ANOVA: *F*_*(1,135)*_ = 3.383, p = 0.07), though there is a significant effect of AM depth on firing rate (*F*_*(1,135)*_ = 5.643, p=0.02) and no interaction between the two variables (*F*_*(1, 135)*_ = 2.013, p = 0.16; Supplemental Figure 3D). However, we found that AC neurons from HL animals displayed weaker temporal coding of AM stimuli. We quantified how well single-unit neurons phase-locked to the AM stimulus by computing the vector strength (VS) at each AM depth (Supplemental Figure 3E) and find that AC neurons from HL animals displayed significantly lower VS values at each depth compared to control neurons (two-way mixed-model ANOVA: *F*_*(1,135)*_ = 5.26, p = 0.023).

Next, we explored whether detection thresholds for the sparse population of individual AC neurons aligned with perceptual performance. When we plot the behavioral threshold as a function of neural threshold, we find a positive correlation for both control and HL animals (Ctrl: Pearson’s *r*: 0.36, p=0.06; HL: *r*: 0.58, p=0.0006; Supplemental Figure 4). This suggests that the neural AM detection thresholds for individual units can explain behavioral performance, but in general underestimates perceptual d’ values. It is therefore likely that animals use more than the information provided by the most sensitive cortical neurons, leading us to explore whether AM detection performance can be better explained by population-level activity.

### An auditory cortex population decoder can explain hearing loss-related behavioral deficits

To determine whether AC population encoding could account for perceptual performance on the AM detection task, we used a previously described procedure (Yao and Sanes, 2018) to construct linear classifiers using support vector machines (SVM) (see Methods). Briefly, AM detection was calculated across our AC neuron population with a linear population readout scheme. The population linear classifiers were trained to decode responses from a proportion of trials to each individual Warn (AM) versus Safe (unmodulated) signal (Figure 6A). Cross-validated classification performance metrics included the proportion of correctly classified Warn trials (“Hits”) and misclassified Safe trials (“False Alarms”). Similar to the psychometric and individual unit neurometric analyses, we converted population decoder performance metrics into d’ values.

**Figure 6.**
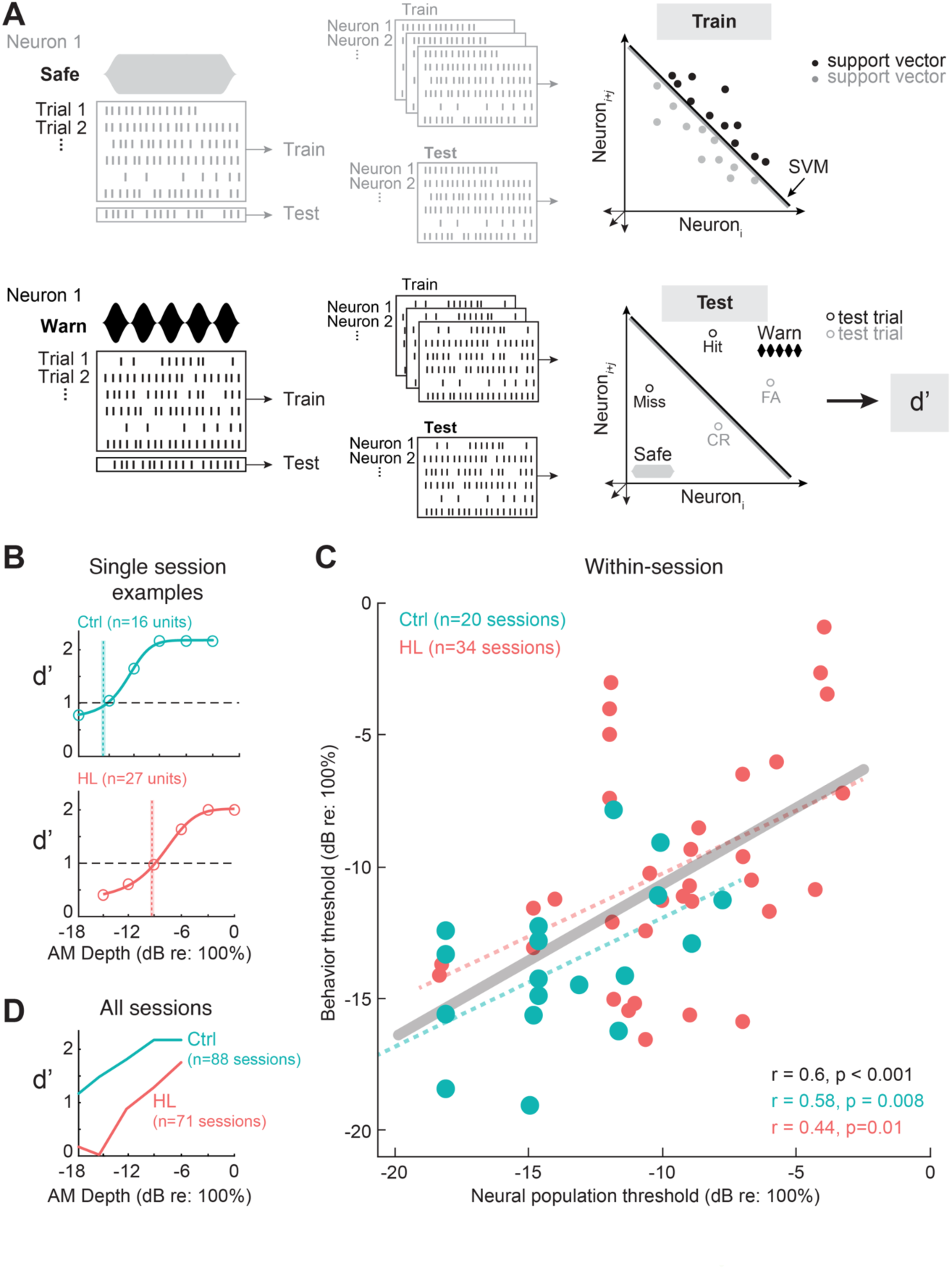
AC population decoder analysis can explain hearing-loss related behavioral deficits. **(A)** Schematic of AC population AM depth encoding with a linear population readout procedure. Hypothetical population responses for individual trials of a Safe (gray; unmodulated) and Warn (black; modulated) stimulus. Spikes were counted across the entire stimulus duration (1 sec) such that spike firing responses from *N* neurons to *T* trials of S stimuli (“Warn” and “Safe”) formed a population “response vector”. A proportion of trials (“leave-one-out” procedure) from each neuron were randomly sampled (without replacement) and fitted to a linear hyperplane that was determined by a support vector machine (SVM) procedure (‘‘train” set). Symbols represent “support vectors”, which are points used to create the linear boundary. Cross-validated classification performance was assessed on the remaining trials (“test” set). Performance metrics included the proportion of correctly classified Warn trials (“Hits”) and misclassified Safe trials (“False Alarms”). Similar to the psychometric and individual unit neurometric analyses, we converted population decoder performance metrics into d’ values. This procedure was conducted across 500 iterations with a new randomly drawn train and test set for each iteration. **(B)** Population decoder performance (d’) for two example individual recording sessions as a function of AM depth from Ctrl and HL neuron populations. Neural d-prime values were fit with a logistic function (solid line), and threshold is defined as the AM depth where the fit crosses d’=1. Sample size indicates the number of single and/or multi-units included within that session. Shaded vertical bars indicate the behavioral threshold for that session. **(C)** Within-session correlations of behavioral threshold plotted as a function of multi-and single-unit population thresholds for that session. Dotted lines indicate the fitted linear regression for Ctrl and HL thresholds, and the solid line indicates the linear regresssion for both groups combined (grey). Pearson’s rand statistical significance of each fit are noted in the bottom right corner of each plot. Behavioral sessions were included if they had at least 10 single and/or multi-units for the decoder analysis. **(D)** Average population decoder performance as a function of AM depth from Ctrl and HL single unit populations pooled across recording sessions.

We applied the population decoder to the data in two ways. First, we assessed decoder performance for single- and multi-units within a behavioral session (i.e., within-session analysis, Figure 6B,C), and second, we assessed decoder performance for single units pooled across all behavioral sessions (Figure 6D). For the within-session analysis, we included sessions that had a minimum of 8 trials per depth recorded from a minimum of 10 multi- and/or single units. Figure 6B shows the population decoder results for an example session from a control and an adolescent HL animal. Similar to the psychometric functions and neurometric data collected for individual single units, the neural d’ values are fit with a sigmoid to calculate neural threshold (where the fit crosses d’=1). Here, the neural thresholds for the example sessions show close alignment with behavioral threshold (vertical bars). When we compared behavioral threshold as a function of neural population threshold for all sessions that meet our criteria (Ctrl: 20 sessions; HL: 34 sessions), we found a strong correlation between behavioral and neural performance (Ctrl: Pearson’s *r*: 0.58, p=0.008; HL: *r*: 0.44, p=0.01; Both groups combined: *r*: 0.6, p<0.001; Figure 6C).

Next, we assessed population decoder performance for single units pooled across behavioral sessions. Units were included in the decoder if behavioral sessions had a minimum number of 8 trials per depth. We opted to restrict the range of depths to −6 to −18 dB re: 100% for both control and HL groups to ensure neurons contributed equally to each depth for both groups. Figure 6D shows the results of the decoder for a population of single units pooled across 85 behavioral sessions (Ctrl: n=34 sessions; HL: 51 sessions) for control and adolescent HL animals (Ctrl: n=5 animals; HL: n=8 animals). Neural thresholds measured from population decoder performance was poorer for single units from adolescent HL animals as compared to control animals (Ctrl: ≤ − 18 dB; HL: −11 dB re: 100%). Furthermore, the neural thresholds measured from population decoder performance closely aligned with perceptual thresholds.

In summary, we find that at the population-level, auditory cortical neurons from adolescent HL animals displayed poorer AM detection thresholds that were closely associated with psychometric performance. This indicates that both individual (Figure 5F) and population-level (Figure 6C,D) activity of AC single units reflect the HL-related AM detection deficits (Figure 3), though with more accuracy at the population-level. Overall, despite over ∼8 weeks of normal audibility following transient adolescent HL, both the neural and behavioral deficit persisted in post-adolescent animals.

### Adolescent hearing loss induced long-lasting changes to auditory cortex excitatory synapses

Transient adolescent HL induced persistent perceptual deficits that were correlated with diminished AC neuron sensitivity, but this effect could have been inherited from lower auditory centers. To test whether poorer neural encoding could be attributed, in part, to the AC, we obtained current clamp recordings in AC thalamocortical brain slices from a subset of behaviorally-tested animals (Ctrl: n=5; HL: n=5).

We first recorded electrical stimulus-evoked inhibitory postsynaptic potentials (IPSPs) from L2/3 pyramidal cells (Ctrl: n=16; HL: n=16) in the presence of ionotropic glutamate receptor antagonists (AP-5; DNQX; Figure 7A,B). Transient adolescent HL did not alter short-latency (putative GABA_A_) IPSP amplitude (one-way ANOVA: *F*_*(1,30)*_ = 0.31, *p* = 0.584; Figure 7C), or long-latency (putative GABA_B_) IPSP amplitude (*F*_*(1,30)*_ = 0.76, *p* = 0.39; Figure 7D). Total IPSP duration was also unaffected by HL (*F*_*(1,30)*_ = 2.76, *p* = 0.11; Figure 7E).

**Figure 7.**
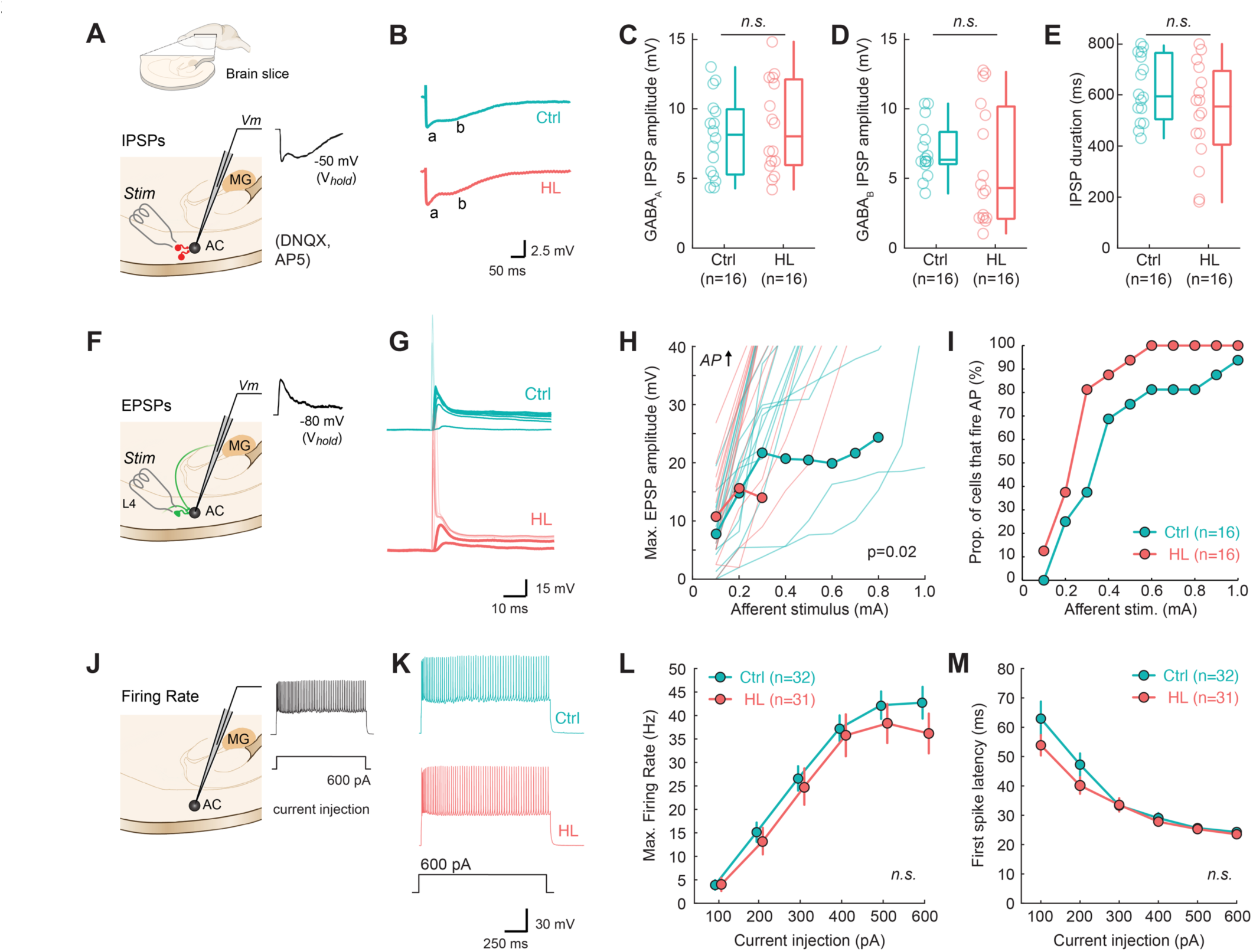
Hearing loss spanning adolescence induces long-lasting changes to auditory cortical properties. **(A)** Schematic of perihorizontal brain slices collected from control (n=5) and HL-reared animals (n=5; inset). Slices contain the auditory cortex (AC), where recordings from layer 2/3 pyramidal cells were made following electrical stimulation (Stim) of fast-spiking inhibitory interneurons to examine synaptic inhibition (IPSPs). Evoked IPSPs were collected in the presence of DNQX and AP-5 to isolate inhibitory potentials. **(B)** Example IPSP traces from control (Ctrl) and HL-reared (HL) cortical cells. Labels a, b indicate putative GABA_A_ and GABA_B_ components, respectively. **(C-E)** GABA_A_ receptor-mediated IPSP amplitudes, GABA_B_ receptor-mediated IPSP amplitudes, and overall IPSP duration are not altered by adolescent HL. (F) Recordings from layer 2/3 pyramidal cells were made following electrical stimulation of local L4 excitatory interneurons to examine synaptic excitation (EPSPs). (G) Example evoked potentials in response to increasing afferent stimulus levels (0.1-1 mA). EPSP traces are in bold and action potentials (APs) elicited are transparent. Stimulus artifact was removed for clarity. (H) Max amplitudes were determined from EPSP waveforms for each stimulation level plotted as input-output functions. Transparent lines indicate individual data, and bold lines with circles indicate mean±SEM if n>3. (I) Proportion of cells that fire APs as a function of afferent stimulation level (mA) for each group. Adolescent HL led to a higher proportion of cells firing APs at lower afferent stimulation levels compared to control cells. **(J)** The firing rate of AC cells were collected via current injection into the cell. **(K)** Example traces of current-evoked responses to a depolarizing current injection (600pA). (L) Input-output function for maximum firing rate in response to current injection steps (100-600pA). HL during adolescence did not significantly impact current-evoked firing rate. **(M)** First spike latency with increasing current injection (pA).

We next recorded electrically-evoked excitatory postsynaptic potentials (EPSPs) from L2/3 pyramidal cells (Ctrl: n=16; HL: n=16) at a holding potential of −80 mV (Figure 7F,G). As shown in Figure 7H, neurons from HL animals required a much lower stimulus current to evoke an action potential (AP). Therefore, a higher proportion of cells from adolescent HL animals fires APs at lower afferent stimulation levels than control cells (Figure 7I), with the average afferent stimulus required to elicit APs of 0.4 mA compared to 0.9 mA in control cells. A two-way mixed model ANOVA revealed a significant effect of HL on EPSP maximum amplitude (*F*_*(1,30)*_ = 5.74, p = .0231), a significant effect of stimulus level (*F*_*(9,22)*_ = 37.52, p<.0001), and no interaction between the two variables (*F*_*(9,22)*_ = 0.853, p = 0.58).Finally, we recorded L2/3 pyramidal cell firing rate in response to current injection (Ctrl: n=32; HL: n=31; Figure 7J,K). Adolescent HL did not alter maximum firing rate (*F*_*(1,61)*_ = 0.649, p = .4274; Figure 7L), or first spike latency (*F*_*(1,21)*_ = 1.21, p=0.29; Figure 7M).

In summary, we find that adolescent HL induces intrinsic and synaptic changes to cortical properties that are distinctly different than the changes observed with critical period HL (Mowery et al., 2015).

### Behavioral and central AM processing deficits were unrelated to peripheral function

One alternative explanation for the AM detection behavioral deficits is that a long duration of HL could induce changes to auditory peripheral structures at, or below, the level of the auditory brainstem. To explore this possibility, we collected auditory brainstem response (ABR) measurements from Control (n=12) and HL (n=13) animals after they completed psychometric testing (Figure 3), and prior to chronic electrode implantation and/or thalamocortical brain slice collection (e.g., see timeline in Figure 1A). Figure 8B shows ABR thresholds in response to a range of tone frequencies (0.5, 1, 2, 4, 5, 8, 16 kHz) for Control and HL animals (i.e., the ABR “audiogram”). The average tone ABR thresholds for HL animals were slightly elevated compared to controls (two-way mixed model ANOVA: *F*_*(1,21)*_ = 8.98, *p* = 0.01). To determine whether this was due to a residual conductive loss or was sensorineural in origin, we examined ABR amplitude input-output functions because subtle damage to auditory nerve synapses has been associated with shallower ABR input-output functions, even when ABR thresholds are normal (Liberman and Kujawa, 2017). Figure 8C shows 4 kHz wave I amplitude (μV) as a function of sound level (dB SPL) for two example ABRs from a control and HL animal. A linear regression was fit to each set of data points (Pearson’s *r*>0.9) which gives rise to a slope value, indicating the wave I amplitude increment of change, or growth, with sound level (μV/dB). For these two examples, the slope is nearly identical (Ctrl: 0.011; HL: 0.012 μV/dB). When we examined all the data collected, we found no differences in the 4 kHz ABR amplitude wave I growth (μV/dB) between control animals (mean ± SEM: 0.0083±0.002 μV/dB) and HL animals (0.0085±0.002 μV/dB). A one-way ANOVA verifies that HL experience did not affect ABR amplitude growth (*F*_*(1,23)*_ = 0.03, *p* = 0.87).

**Figure 8.**
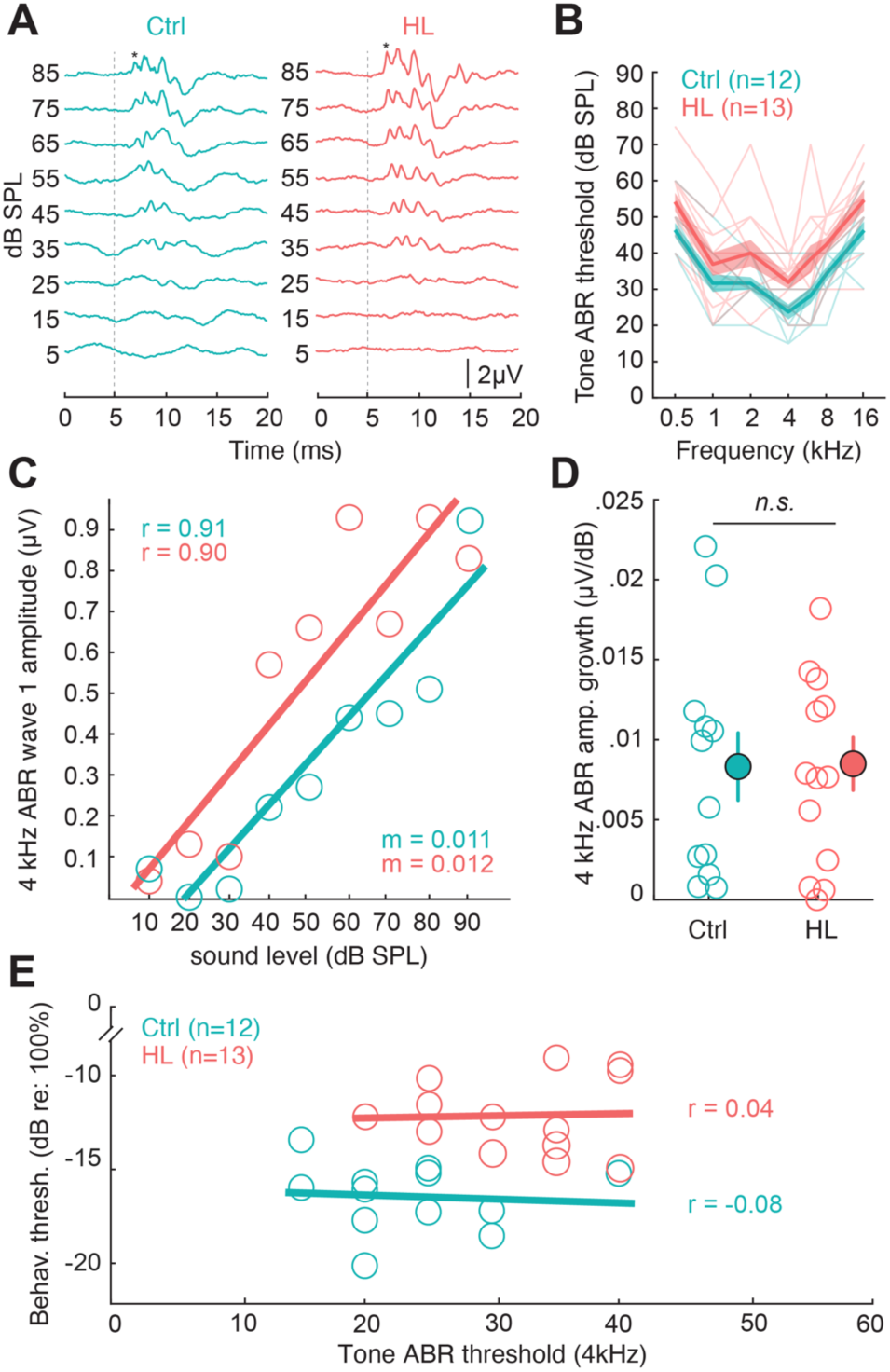
Control experiment: HL-related AM detection deficits are not due to status of auditory periphery. **(A)** Auditory brainstem response (ABR) waveforms in response to 4kHz tone pips for example control (Ctrl, Subject ID: M274129) and HL-reared animal (HL, Subject ID: M274131). Vertical dotted line indicates stimulus onset. Asterisk (*) indicates ABR wave 1. **(B)** Tone ABR Thresholds (mean±SEM) as a function of frequency (kHz). **(C)** 4kHz ABR wave 1 amplitude (μV) as a function of sound level (dB SPL) for two example Ctrl and HL animals. Solid line indicates linear fit; m indicates the slope of the fit. Pearson’s *r* value for each regression are noted on the top left corner. **(D)** 4kHz ABR wave 1 amplitude growth is comparable between Ctrl and HL-reared animals. Amplitude growth values are calculated from the slope of the linear regression to wave 1 amplitude with sound level (i.e., in C). **(E)** Individual behavioral threshold as a function of ABR threshold at 4 kHz. Behavioral thresholds were from the final day of psychometric testing, closest to when the ABR was collected. Solid line indicates fitted linear regression. Horizontal linear fits indicate no correlation (Pearson’s r ≤0.04 for both groups) between tone ABR thresholds and behavioral thresholds.

Finally, if ABR thresholds were causally related to behavioral thresholds, then we would expect a correlation between them. Figure 8E plots individual behavioral thresholds as a function of 4 kHz ABR threshold for Control and adolescent HL animals. Fitted linear regressions displayed no correlation (Ctrl, Pearson’s *r* = −0.08; HL, *r* = 0.04).

Taken together, Control and HL animals both exhibit comparable ABR wave I amplitude growth with sound level indicating that the auditory nerve fibers were not compromised following adolescent HL. Moreover, we found no relationship between tone ABR thresholds and behavioral performance, suggesting that the perceptual deficit we observed was due to central changes at the level of the auditory cortex, not peripheral changes at, or below, the level of the auditory brainstem.

### AM depth detection thresholds remained stable across relevant sound levels

To further test the possibility that a residual conductive loss could contribute to AM detection deficits, we determined how behavioral depth detection thresholds vary with sound level. If thresholds do not vary across a large range of sound levels, then that would suggest a small residual conductive loss could not explain the behavioral findings.

A group of normal hearing animals (n=4) were trained on the AM depth detection task (Figure 9A). After obtaining AM detection thresholds at the same sound level used for the experimental groups (45 dB SPL; Figure 9B), the sound level was varied from 10-60 dB SPL across 24 testing sessions, with at least 3 sessions per level (Figure 9C). Figure 9D shows depth detection thresholds as a function of sound level for all animals. A one-way ANOVA revealed a significant effect of sound level on detection thresholds (*F*_*(7,87)*_ = 3.25, *p* = 0.004), and a Tukey HSD *post hoc* test for pairwise comparisons revealed 10 dB SPL was significantly different from all other levels (p<0.0001), and 15 dB SPL was significantly different from 40 dB SPL (*t* = 3.21, *p* = 0.04), 45 dB SPL (*t* = 3.92, *p* = 0.004), and 60 dB SPL (*t* = 3.73, *p* = 0.008). All other sound levels were not significantly different from one another (p=0.3-1.0). To determine the level at which thresholds remained consistent with sound level, a 3-parameter exponential was fit to the data (dotted line in Figure 9D). We found that thresholds asymptote at −15.8 dB (re: 100% AM), which corresponds to 27 dB SPL. This indicates that depth detection thresholds remain stable for sound levels greater than 27 dB SPL, which is well below the sound level used in the behavioral and awake-behaving experiments (45-60 dB SPL). This suggests that even if an animal has a mild peripheral HL (<20 dB), behavioral detection thresholds would not be significantly impacted. This further supports the notion that the behavioral deficit we observed in animals after adolescent HL is due to central changes at the level of the AC, and not due to peripheral auditory mechanisms.

**Figure 9.**
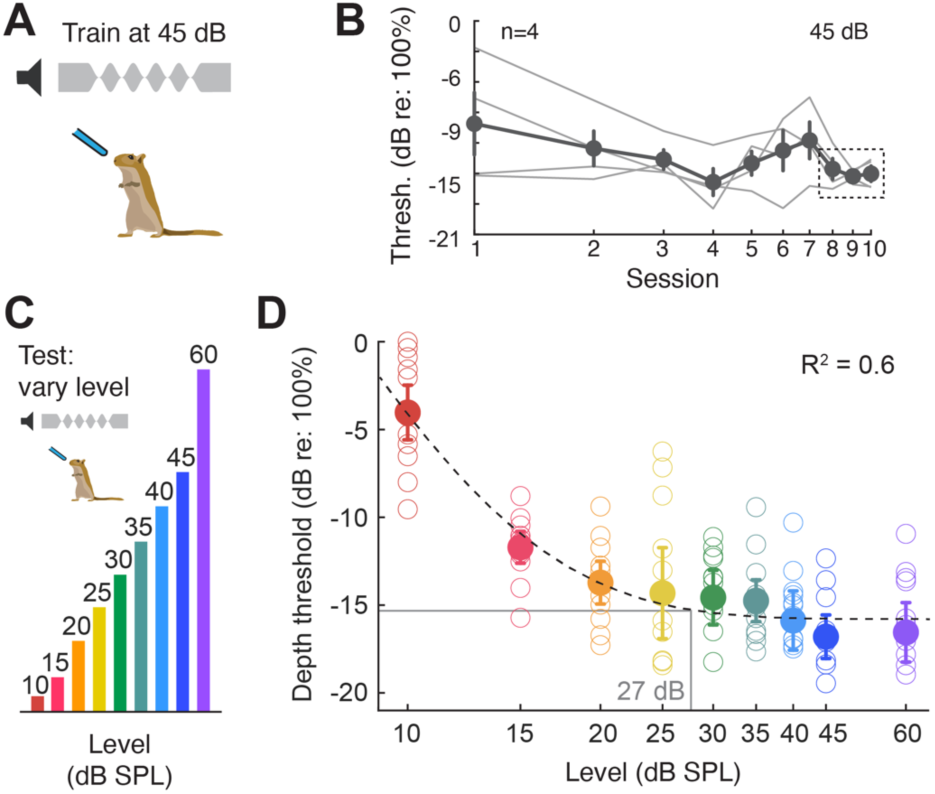
Control experiment: AM depth detection thresholds remain consistent with varying sound levels. **(A)** Animals (n=4) were trained on the AM depth detection task at 45 dB SPL. **(B)** Depth detection thresholds as a function of 10 testing days. Once detection thresholds stabilize (i.e., consistent for 3 consecutive sessions: dotted rectangle), then the sound level was varied from 10-60 dB SPL **(C). (D)** Behavioral AM depth detection thresholds across sound level (dB SPL). Open circles indicate individual performance (3 sessions per animal) and solid circles indicate the group mean (± SEM). A 3-parameter exponential was fit to the data (dotted line). Thresholds asymptote at −15.8 dB (re: 100% AM), which corresponds to 27 dB SPL. Therefore, depth detection thresholds remain consistent for sound levels greater than 27 dB SPL.

## Discussion

The extent to which perceptual skills are influenced by adolescent sensory experience remains uncertain. Before sexual maturity, the influence of sensory experience is profound, and brief periods of diminished or augmented stimulation can permanently alter central nervous system function (Hubel and Wiesel, 1970; Van der Loos and Woolsey, 1973; Knudsen et al., 1984b; Hensch, 2005; de Villers-Sidani et al., 2007; Han et al., 2007; Zhou and Merzenich, 2009; Popescu and Polley, 2010; Christakis et al., 2012; Cheetham and Belluscio, 2014; Mowery et al., 2015). However, both CNS function and perception continue to mature after the onset of sexual maturation (Carey et al., 1980; Sharma et al., 1997; Giedd et al., 1999; Moore and Guan, 2001; Bishop et al., 2007; Sussman et al., 2008; Guyer et al., 2008; Pinto et al., 2010; Banai et al., 2011; Huyck and Wright, 2013; Lebel and Beaulieu, 2011; Petanjek et al., 2011; Germine et al., 2011; Huyck and Wright, 2011; Mahajan and McArthur, 2012b, 2012a; Kadosh et al., 2013; Selemon, 2013; Cohen Kadosh et al., 2013; Dayanidhi et al., 2013; Shafer et al., 2015; Simmonds et al., 2017; Downes et al., 2017; McMurray et al., 2018), suggesting that sensory plasticity could remain heightened during adolescence. Here, we report that a period of mild auditory deprivation during adolescence, but not adulthood, induced long-lasting perceptual deficits that could be explained by intrinsic changes to auditory cortex (AC) excitatory synapse function and degraded AC population encoding during task performance.

### Comparison between juvenile and adolescent plasticity

Mammals display enhanced plasticity from early infancy through juvenile development, allowing for both positive and negative environmental influences. Foremost among these is the early influence of sensory experience: finite periods of monocular deprivation shortly after eye opening result in long-term visual acuity deficits that are closely associated with degraded cortical processing (Hubel and Wiesel, 1970). Early critical periods are found in humans: children born with dense cataracts exhibit long-term visual acuity deficits even after cataract removal (Lewis and Maurer, 2005; Jain et al., 2010) and, similarly, those born with profound hearing loss display a neonatal epoch during which cochlear prostheses provide maximal restoration of neural and behavioral performance (Sharma et al., 2002).

Adolescent plasticity displays some unique characteristics, as neural function and behavioral skills slowly transition to an adult phenotype (Sawyer et al., 2018). For example, there is a transient reduction in some forms of learning or memory. This includes slower rates of auditory perceptual learning (Huyck and Wright, 2011; Caras and Sanes, 2019), reduced voice or face recognition (Mann et al., 1979; Carey et al., 1980), and diminished extinction learning (McCallum et al., 2010; Kim et al., 2011; Pattwell et al., 2012). There are also indications that adolescents are vulnerable to environmental factors. For instance, social experience during early adolescence is particularly important for the subsequent development of these skills (Burke et al., 2017). Rats that experience social isolation from a late juvenile to early adolescent age exhibit abnormal exploratory behavior when placed in a novel environment (Einon and Morgan, 1977), while social deprivation before or after this age range is not damaging. In addition, rats that experience chronic stress throughout adolescent development exhibit long lasting structural changes in the hippocampus that coincide with spatial navigation impairments (Isgor et al., 2004). Our results suggest that adolescent vulnerability noted above extends to skill learning. We found that transient adolescent HL led to significantly poorer perceptual learning of the auditory task (Figure 3C).

### Effects of adolescent sensory deprivation

The current findings suggest that adolescent neural plasticity differs dramatically from that observed during the AC critical period. One novel outcome from this study is that critical period and adolescent HL induce similar behavioral deficits, but each are associated with unique synaptic mechanisms. Transient HL during the critical period (P11-23) causes reduced inhibitory synaptic strength and a decrease in membrane excitability in AC neurons (Kotak et al., 2005, 2008; Takesian et al., 2010, 2012; Mowery et al., 2015, 2019). In contrast, we found that transient HL spanning adolescence had no effect on synaptic inhibition or current-evoked firing rates. Rather, adolescent HL led to elevated excitatory postsynaptic potential amplitudes in AC pyramidal neurons (Figure 7). A second novel outcome is that adolescent HL leads to a significant deterioration of AC stimulus encoding during task performance (Figures 5 and 6), despite having no effect on neural discharge rate (Supplemental Figure 3). This is in marked contrast to critical period HL which causes a significant decrease of AC neuron firing rate (Rosen et al., 2012; Yao and Sanes, 2018). Taken together, these findings suggest that adolescent neural plasticity in response to sensory deprivation is mechanistically distinct from that observed during early critical periods.

A third novel finding is that the effects of adolescent HL are more persistent than those observed following brief HL during the AC critical period. For example, restoring normal auditory input at P17 in the gerbil, before closure of the AC critical period (P23), led to a near-complete recovery of intrinsic cellular properties whereas restoring auditory input after the critical period ends (P23) led to persistent AC alterations (Mowery et al., 2015). A transient period of developmental HL can also cause perceptual deficits, though these often resolve gradually (Hall et al., 1995; Moore et al., 1999; Caras and Sanes, 2015; McKenna Benoit et al., 2018). Thus, we previously reported that brief HL during an AC critical period (P11-23) led to a perceptual deficit, while the same manipulation had no effect when initiated after the critical period closed (P23) (Caras and Sanes, 2015). In contrast, our current study found significant and persistent perceptual impairments following HL that was induced at the same postnatal age, P23. This suggests that auditory function is susceptible to deprivation after the AC critical period, but only if the deprivation persists throughout adolescence. In fact, we found that adolescent HL induced longer lasting perceptual deficits than those observed with critical period HL. Following adolescent HL, neural and behavioral deficits were still present for over ∼8 weeks following restoration of normal auditory input, whereas the critical period-induced perceptual deficits largely resolved over the same recovery period (Caras and Sanes, 2015).

### Alternative explanations for the effect of auditory deprivation

We find that adolescent HL induces intrinsic changes to AC neurons that contributes to poorer AM detection thresholds long after normal hearing is restored. However, it is also possible that the long duration of the hearing deprivation compromised peripheral auditory structures, resulting in decreased audibility. For instance, subtle damage to auditory nerve synapses is evidenced by shallower ABR input-output functions even with normal ABR thresholds (Furman et al., 2013; Liberman and Kujawa, 2017). However, our ABR measurements reveal no significant differences in ABR wave I rate of amplitude growth with sound level (Figure 8D), indicating intact auditory nerve function. Although we find a modest difference in tone-ABR thresholds following adolescent HL, this does not account for poorer perceptual performance (Figure 8E). Furthermore, we find that AM detection thresholds are largely independent of sound level (above 27 dB), consistent with other assessments of AM depth sensitivity (Viemeister, 1979). This suggests that even if an animal has a mild residual peripheral HL, AM detection thresholds would not be significantly impacted as the sound level used in the behavioral and awake-behaving experiments far exceed the potential HL. In addition, permanent conductive HL (∼35-40 dB) does not appreciably alter frequency tuning in the cochlea, suggesting that associated perceptual dysfunction is due to central mechanisms (Ye et al. (2021). Although we did not examine AM-following responses (i.e., an assessment of brainstem temporal processing) in the current study, we previously found that permanent conductive HL does not alter this property, despite causing perceptual and cortical deficits (Yao and Sanes, 2018).

A second possibility is that the adolescence, itself, was disrupted by the manipulation. To directly identify the period of adolescence in the gerbil, and confirm that it was not altered by our manipulation, we measured testosterone levels from each subject throughout development. We identified a developmental window during which testosterone levels increased, and found it to be identical between normal hearing and earplugged animals, and consistent with a previous report (Pinto-Fochi et al., 2016). Since female gerbils generally reach sexual maturity concurrently with or prior to their male counterparts (Norris and Adams, 1974), it is plausible that the HL manipulation spanned sexual development for both sexes. Furthermore, we found estradiol levels to be comparable between normal hearing and earplugged animals at P90-102, indicating that the manipulation did not alter estradiol levels by the time adulthood was reached. Therefore, a change to adolescent maturation was unlikely to account for our results. In addition, sex hormones can give rise to sex-specific differences in auditory processing and perceptual skills (e.g., Canlon and Frisina, 2009). However, we did not find sex-specific differences in procedural learning or psychometric task performance between male and female control or adolescent HL gerbils (Supplemental Figure 1). It was also possible that the earplug manipulation caused chronic stress during adolescence which adversely effects the CNS (Isgor et al., 2004). However, cortisol levels did not differ between treatment groups (Supplemental Figure 2), and displayed no correlation with perceptual performance. Therefore, chronic stress was unlikely to explain our findings.

Another potential explanation for our behavioral findings is that adolescent HL may influence the functional properties of neurons downstream from AC that mediate cognitive processing. In fact, transient HL during the auditory cortex critical period causes persistent change to auditory striatal neurons (Mowery et al., 2017). Areas such as the prefrontal cortex that support cognitive and executive function continue to maturate into young adulthood, and may remain vulnerable to deprivation (Kolb et al., 2012). Furthermore, HL is thought to be a risk factor for cognitive dysfunction in humans: Childhood HL can delay maturation of cognitive skills such as auditory working memory and attention (Reichman and Healey, 1983; Pisoni and Cleary, 2003; Holmes et al., 2017; Moore et al., 2020). Thus, it is possible that HL may influence the development of both sensory and non-sensory circuits.

### The timing and duration of developmental hearing loss

Sensory manipulations that target adolescence are somewhat uncommon. In the auditory system, manipulations have primarily targeted either early development, shortly after hearing onset, or adults, well after sexual maturity. For instance, transient HL (i.e., using earplugs, ear canal ligation, or poloxamer hydrogel) has been induced at hearing onset in the guinea pig (Clements and Kelly, 1978), gerbil (Caras and Sanes, 2015; Mowery et al., 2015, 2017, 2019; Green et al., 2017), barn owl (Knudsen et al., 1982, 1984b, 1984a; Knudsen, 1983, 1985; Mogdans and Knudsen, 1992, 1993), ferret (Moore et al., 1999; Keating et al., 2013, 2015a), rat (Popescu and Polley, 2010), and mouse (Polley et al., 2013; Zhuang et al., 2017). Permanent forms of mild-moderate HL (i.e., conductive HL through ossicle disruption or surgical intervention of the ear canal) have also been induced at hearing onset of a variety of species (gerbil: Xu et al., 2007; Rosen et al., 2012; Takesian et al., 2012; Buran et al., 2014a; Gay et al., 2014; Ihlefeld et al., 2016; von Trapp et al., 2017; Yao and Sanes, 2018; rat: Clopton and Silverman, 1977; Silverman and Clopton, 1977; mouse: Xu and Jen, 2001; cat: Moore and Irvine, 1981; Brugge et al., 1985).

The duration of developmental HL manipulations vary widely, as does its relationship to the age of sexual maturity. For instance, Polley et. al. (2013) induced a ∼7 day auditory deprivation beginning at P12 in the mouse, a species that reaches sexual maturity around postnatal day 49 (Danneman et al., 2012). A similar duration of HL would represent a much smaller fraction of development for species with longer periods of adolescence (e.g., sexual maturity in the ferret is 4-6 months or ∼1 year in the barn owl). Though the majority of transient HL studies focus on relatively brief deprivation early in development, there are a few that span the late juvenile through adolescent period. A reversible HL was induced in barn owls beginning at posthatch day 28, approximately 20 days after hearing onset, and normal audibility was restored between days 120-213, prior to the age of sexual maturity at ∼1 year (Knudsen et al., 1984a; Knudsen et al., 1985; Mogdans and Knudsen, 1992; Mogdans and Knudsen, 1993). Similarly, a transient HL was induced in the ferret at P24-25, just before hearing onset, and audibility was restored at P126 or P213, after sexual maturity at ∼P121-152 (Moore et al., 1999; Keating et al., 2013; Keating et al., 2015b). The relatively long duration developmental HL in both the barn owl and ferret studies induced long-lasting neural and behavioral deficits, similar to our current findings. However, none of the previously reported studies exclusively targeted adolescence after the critical period.

Our results are also consistent with clinical evidence that suggests longer durations of developmental HL can induce more harmful effects. Children with progressive, late-onset, or acquired HL (Smith et al., 2005) often pass newborn hearing screenings, leading to long periods of undetected HL. Those with an extended period of unresolved HL display more severe language deficits than those with a brief HL (Yoshinaga-Itano et al., 1998; Tomblin et al., 2015). Here, we find that a long duration HL that spans adolescence disrupts perceptual performance on an auditory detection task, whereas HL of the same duration induced in adulthood did not alter detection performance. While we did not observe deficits following adult-onset HL, it is possible that long duration HL would be detrimental if induced in older adult animals as HL in aging humans is linked with deficits in cognitive and central auditory processing (Henkin et al., 2014; Finke et al., 2015) that pose a greater risk for cognitive decline and dementia if left untreated (Lin et al., 2013). We then found that long duration HL that spans adolescence led to AC neural changes that are distinctive from critical-period plasticity during juvenile development. We conclude that the changes to AC synaptic excitability observed following adolescent HL contributes to the AC encoding deficits observed with the awake behaving neural recordings (Figures 5-6). Taken together, these results reveal that transient HL spanning adolescence alters AC cellular properties long-term, thereby diminishing sensory encoding of AM stimuli during the detection task, and explains the long-term auditory perceptual limitations observed in adolescent HL animals.

## Methods

### Experimental subjects

A total of 46 Mongolian gerbils (*Meriones unguiculatus*, 26 females, 20 males) were used in the study. Pups were weaned from commercial breeding pairs (Charles River Laboratories) at postnatal (P) day 30 and housed on a 12-hour light / 12-hour dark cycle with full access to food and water, unless otherwise noted. All procedures were approved by the Institutional Animal Care and Use Committee at New York University.

### Hormone assessment

Estradiol, testosterone, and cortisol hormone levels were collected in developing gerbils through blood sample collection at five timepoints spanning development (P35, P55, P65, P90, and P102). Animals were briefly anesthetized with isoflurane/O2 and placed on a heating pad. The fur at the base of the tail was removed using a scalpel blade for greater visualization and access to the lateral tail vein. A small incision was made and the blood from the lateral tail vein was collected in Eppendorf vials (100-300μl). To reduce acute stress-induced changes to cortisol levels, all blood collection was obtained within 3 minutes of placing animals under anesthesia. Blood collection was also collected at the same time of the day for each animal at each testing age (early afternoon) to avoid circadian-related hormonal fluctuations. Blood samples were centrifuged (for 9 minutes, 8,000g) within 30 minutes to avoid hemolysis of the blood. After spinning, the serum component of the blood was extracted into a separate vial and stored in a − 80° freezer until samples were ready to be processed.

Serum samples were shipped to the Endocrine Technologies Core (ETC) at the Oregon National Primate Research Center (ONPRC) for quantification of hormone levels. Serum concentrations of estradiol (E2), testosterone (T), and cortisol (F) were simultaneously measured by liquid chromatography-tandem triple quadrupole mass spectrometry (LC-MS/MS) using a previously established method(Bishop et al., 2019). Briefly, serum samples were combined with a mixture of deuterium-labeled internal standards for E2, T, and F. Steroids were extracted using supported liquid extraction (SLE) and analyzed on a Shimadzu Nexera-LCMS-8050 LC-MS/MS platform. 9-point calibration curves for each hormone were linear throughout the assay range (13 pg/ml – 26.67 ng/ml; R>0.99). The lower limit of quantitation for each hormone was 13 pg/ml. Intra-assay CVs were 15% for E2, 2.9% for T, and 3.8% for F. Serum testosterone and cortisol levels were successfully obtained using the LC-MS/MS approach; however, estradiol was not detected in any of the serum samples indicating that the assay was not sensitive enough to determine estradiol levels (i.e., values were below the lowest calibration level and could not be quantified). To quantify estradiol levels, the remaining serum samples for the P90 and P102 timepoints were then pooled to be tested on a more sensitive LC/MS system. The serum samples were pooled as follows: (1) Control, Male, age: P90-P102 (12 total samples; 5 subjects), (2) Control, Female, age: P90-102 (11 total samples; 7 subjects), (3) Hearing loss (P23-102), Male, age: P90-102 (12 total samples; 5 subjects), (4) Hearing loss (P23-102), Female, age: P90-102 (16 samples; 9 subjects). The pooled samples were then sent to the Endocrine Technologies Core (ETC) at the Oregon National Primate Research Center (ONPRC) for quantification of estradiol using a more sensitive LC/MS instrument, the Shimadzu Nexera-LCMS-8060 LC-MS/MS, where the lower limit of quantification for estradiol was 3 pg/ml.

### Transient auditory deprivation

Custom earplugs (BlueStik adhesive putty, super glue, and KwikSil silicone adhesive) were used to induce a reversible, transient hearing loss (HL). Earplugs provided a mild-moderate temporary HL, attenuating ∼10-20 dB at frequencies ≤ 1 kHz and 30-50 dB at frequencies ≥2 kHz (Caras and Sanes, 2015). To assess vulnerability in adolescence, gerbils either received bilateral earplugs from P23 (after the auditory cortex critical period) through P102 (after sexual maturity; n=14) or no earplugs (littermate controls, n=12). To assess vulnerability in adulthood, gerbils either received bilateral earplugs beginning ≥P102 (after sexual maturity; n=8) for the same duration as the adolescent HL animals (80 days) or no earplugs (littermate controls, n=8). Animals were briefly placed under isoflurane anesthesia and the ear canal and tympanic membrane were visualized under a stereomicroscope (Olympus). A small adhesive putty “disk” was gently inserted into the ear canal and compressed into the bony portion of the canal. Care was taken to ensure that the earplug material did not go near the tympanic membrane. A small drop of super glue was placed onto the putty and allowed to fully dry before covering the earplug with a layer of KwikSil silicone adhesive. Earplugs were checked 1-2x daily and replaced as needed. Following earplug removal, the tympanic membrane was visualized (at the time of removal as well as post-mortem) to ensure the membrane was intact and free from debris. Littermate control animals also underwent daily handling and had comparable isoflurane exposure with similar manipulations of their pinna (i.e., used forceps to mimic earplug placement) as their earplugged counterparts. Animals were allowed to recover for 21 days prior to behavioral and physiological testing.

### Behavioral training and testing

#### Behavioral apparatus

All behavioral experiments were performed in a sound-attenuating booth (Industrial Acoustics) and were observed through a closed-circuit video monitor (Logitech HD C270 Webcam). Animals were placed in a custom-made, acoustically-transparent plastic test cage with a stainless steel water spout positioned above a metal platform. Sound stimuli were delivered from a calibrated free-field speaker (DX25TG0504; Vifa) positioned 1 m above the test cage. Contact with the waterspout was assessed via infrared sensor (OP550 NPN Silicone phototransistor, Digikey) and emitter (940 nm; LTE-302, Digikey) housed within a custom apparatus encasing the waterspout. Acoustic stimuli, water reward delivery, experimental parameters, and data acquisition were controlled using custom MATLAB scripts developed by Dr. Daniel Stolzberg (https://github.com/dstolz/epsych) and using a multifunction processor (RZ6; Tucker-Davis Technologies).

#### Procedural training

Animals were trained on an amplitude modulation (AM) depth detection task using an aversive conditioning Go/Nogo procedure used in the laboratory (Sarro and Sanes, 2010; Sarro et al., 2011; Sarro and Sanes, 2011; Rosen et al., 2012; Buran et al., 2014a; Caras and Sanes, 2015, 2017). Gerbils were placed on controlled water access and trained to drink from a waterspout in the presence of continuous unmodulated noise (60 dB SPL; band-limited white noise, 3-20 kHz; Nogo/”Safe” stimulus), and to refrain from drinking when the unmodulated noise transitioned to a 5 Hz AM rate signal (Go/”Warn” stimulus). This was done by pairing the AM signal with a mildly aversive shock (300 ms; 0.5-1.0 mA; Lafayette Instruments) delivered through the waterspout. Modulated, “warn” trials (1 sec AM stimulus) were randomly interspersed with 3-5 unmodulated, “safe” trials (1 sec trial duration) to deter temporal conditioning. First, animals were trained on the most salient AM cue (100% modulated depth). Behavioral responses were classified as a “hit” if the animal correctly withdrew from waterspout during modulated warn trials and as a “false alarm” if the animal incorrectly withdrew from waterspout during unmodulated safe trials (see schematic in Figure 2A). Behavioral performance was quantified using the signal detection metric, *d’*, defined as *d’ = z(hit rate) – z(false alarm rate)* (Green and Swets, 1966). Animals achieved task proficiency when *d’* ≥ 1.5 and completed three separate training sessions with a sufficient number of warn trials (≥ 50). On the final session of procedural training, the sound level was lowered to 45 dB SPL.

#### Perceptual testing

Once animals reached the training performance criterion (*d’* ≥ 1.5), animals were then perceptually tested on a range of AM depths, from fully modulated (100% AM) to shallow modulations (> 0% AM). Note, AM depths were presented on a dB scale (re: 100% depth), where 0 dB (re: 100% AM) refers to fully modulated noise (100%) and decreasing (negative) values refer to smaller depths. These values are not to be confused with overall stimulus intensity, which remained at 45 dB SPL during perceptual testing. Furthermore, the gain of the AM stimuli was adjusted to accommodate for changes in average power across modulation depth (Wakefield and Viemeister, 1990). Perceptual performance was assessed over ≥10 testing sessions. For each session, the range of AM depths presented was adjusted to bracket the animal’s detection threshold, which was defined as the AM depth at which *d’=1*.

#### Psychometric analysis

Classified behavioral responses (i.e., Hits, False alarms) were used to generate psychometric functions used to determine AM depth detection thresholds. First, psychometric functions based on the percentage of Hits were plotted as a function of AM depth and fit with cumulative Gaussian using Bayesian inference (open-source MATLAB package psignifit 4) (Schütt et al., 2016; Caras and Sanes, 2017; Yao and Sanes, 2018). The default priors in psignifit 4 were used as it fit the behavioral data with high accuracy. Psychometric fits were then transformed to *d’* values (*d’ = z(hit rate) – z(false alarm rate)*(Green and Swets, 1966). Psychometric functions using d’ were used to determine AM depth detection thresholds, which was defined as the AM depth at which *d’=1*. When computing *d’*, we constrained the hit and false alarm rates (floor: 0.05; ceiling: 0.95) to avoid *d’* values that approached infinity (Caras and Sanes, 2017; Yao and Sanes, 2018).

### In vivo neurophysiology

#### Electrode implantation

The procedural details for electrode implantation were identical to the procedures used in previous studies from our laboratory (Buran et al., 2014b; von Trapp et al., 2016; Caras and Sanes, 2017; Yao and Sanes, 2018). The procedures are briefly described below.

Following behavioral assessment, a subset of animals (Control: n=7; HL: n=9) were implanted with a multichannel electrode array in the primary auditory cortex (AC). Animals were anesthetized with isoflurane/O2 and secured on a stereotaxic apparatus (Kopf). The skull was exposed and a craniotomy was made on the dorsal skull at established coordinates for targeting the left core AC. The left cortex was targeted as it is more responsive to temporal acoustic features (Heffner and Heffner, 1984) A 64 channel silicone probe array (Neuronexus: Buzsaki64-5×12_H64LP_30mm or A4×16-Poly2-5mm-23s-200-177-H64LP_30mm) was implanted into the AC and oriented such that the array spanned the tonotopic axis (Ter-Mikaelian et al., 2007; Rosen et al., 2012; von Trapp et al., 2016). The array was positioned at a 25º angle in the mediolateral plane, and the rostral-most probe shank was positioned at 3.9 mm rostral and 4.6-4.9 mm lateral to lambda. The array was fixed to a custom-made microdrive to allow for later advancement of the electrode across multiple behavioral testing days. Following recovery, neural recordings were made while animals performed the AM detection task. When all testing was complete, animals were deeply anesthetized, perfused with fixative (phosphate-buffered saline, 4% paraformaldehyde), brain extracted, sectioned (Leica vibratome), and stained with Nissl dye. Electrode track position in the core AC was verified using a gerbil brain atlas (Radtke-Schuller et al., 2016).

#### Data acquisition

Extracellular recordings were acquired using a wireless recording system (Triangle BioSystems) while animals performed the AM detection task. Signals from all channels were pre-amplified, digitized (24.4 kHz; PZ5; Tucker Davis Technologies, TDT), and sent to the RZ2 (TDT) for filtering and processing. All recordings were pre-processed and sorted offline. First, recordings were pre-processed using custom MATLAB scripts and initial spike thresholding was performed. Spikes were semi-automatically sorted using Kilosort, a spike sorting framework based on template matching of spike waveforms (Pachitariu et al., 2016). Spike waveforms were manually inspected and refined in Phy (Rossant et al., 2016). Units were classified as single units if they were well-isolated, displayed clear separation in PC space, and had few refractory period violations (<10%). Units that did not meet criteria for single unit classification were classified as multi-units. Sorted spike data were then analyzed using custom MATLAB scripts.

#### Neurometric analysis

Procedures for assessment of neural AM detection are similar to that described previously in the laboratory (Caras and Sanes, 2017). Briefly, the firing rate (spikes / second) for each unit was calculated over a 1 second duration for modulated and unmodulated stimuli. The 1 sec time period corresponds to the duration of each trial. Neural *d’* values were computed based on the firing rate, where for each AM depth, the firing rate was normalized by the standard deviation pooled across all the stimuli (i.e., z-score), and subtracting the unmodulated value from each AM stimulus (*d’*_FR_= z(AM Depth) - z(Unmodulated)).

The neural *d’* values were used to generate neurometric functions, where the data were fit with a logistic function using a nonlinear least-squares regression procedure (MATLAB function *nlinfit*). See Supplementary Figure 3C for fit of neural d’ data for two example neurometric functions. The neurometric fit was deemed appropriate if the correlation between the predicted and actual d’ values were statistically significant (Pearson’s *r*). As with the perceptual definition, AM detection thresholds for each unit were defined as the AM depth at which *d’*=1. Units were classified as AM responsive if (1) the neurometric fit was significant and (2) the maximum neural *d’* value was ≥1. All other units that did not meet the criteria for AM responsivity were classified as unresponsive.

In addition, the strength of synchronization of unit responses to the envelope of each AM stimulus was quantified using a vector strength (VS) metric (Goldberg and Brown, 1969). Spikes occurring at the same phase of the AM resulted in high synchrony (VS=1) whereas spikes out of phase resulted in low synchrony (VS=0). The Rayleigh test of uniformity was used to determine statistical significance of the VS (Mardia, 1972) at the level of p<0.001.

#### Population coding

We used a previously employed linear classifier readout procedure (Yao and Sanes, 2018) to assess AM detection across a population of AC neurons. A linear classifier was trained to decode responses from a proportion of trials to each stimulus set (e.g., “Warn” and “Safe”; Figure 6A). Specifically, spike count responses from all neurons were counted across 1000 ms from “T” trials of “S” stimuli (e.g., AM depth) and formed the population “response vector”. Since the number of trials were unequal across all units, we randomly subsampled a proportion of trials (i.e., 8 trials) from each unit. 7 of the 8 trials were then randomly sampled (without replacement) across all neurons. A support vector machine (SVM) procedure was used to fit a linear hyperplane to the data set (“training set”). Cross-validated classification performance was assessed on the remaining single trial (1 of the 8 trials) by computing the number of times this test set was correctly classified and misclassified based on the linear hyperplane across 500 iterations with a new randomly drawn sampled train and test sets for each iteration. Performance metrics were computed to determine the “Hit” rate (proportion of correctly classified Warn trials) and “False Alarm” rate (proportion of misclassified Safe trials). Similar to the psychometric and individual unit neurometric analyses, we converted population decoder performance metrics into d’ values. The SVM procedure was implemented in MATLAB using the “fitcsvm” and “predict” functions with the “KernelFunction” set to “linear”.

#### Histology

After all electrophysiology experiments, animals were given an overdose of sodium pentobarbital (150 mg/kg, intraperitoneal injection) and perfused with 0.01 M phosphate-buffered saline followed by 4% paraformaldehyde. The brains were extracted, post-fixed in 4% paraformaldehyde, and embedded in 6% agar and sectioned at 60 μm on a vibratome (Lieca). Tissue sections were mounted on gelatin-subbed glass slides, dried, and stained for Nissl. Brightfield images were acquired using an upright microscope (Revolve, Echo) and electrode tracks were reconstructed offline and compared with a gerbil brain atlas (Radtke-Schuller et al., 2016) to confirm the electrodes targeted core auditory cortex (Figure 5B).

### In vitro electrophysiology

#### Thalamocortical brain slice preparation

Following behavioral data collection, a subset of animals (Control: n=5; HL: n=5) were used for *in vitro* auditory cortex (AC) recordings. The procedures for brain slice physiology experiments have been established in prior studies from our laboratory (Kotak, 2005; Xu et al., 2007; Takesian et al., 2012; Mowery et al., 2015). Animals were deeply anesthetized (400 mg/kg, chloral hydrate, intraperitoneal) and brains extracted and placed into 4ºC oxygenated artificial cerebrospinal fluid (ACSF, in mM: 125 NaCl, 4 KCl, 1.2 KH2PO4, 1.3 MgSO4, 26 NaHCO3, 15 glucose, 2.4 CaCl2, and 0.4 L-ascorbic acid, pH=7.4 after bubbling with 95% O2/5% CO2). Brains were sectioned horizontally (500 μm) to preserve the ventral medial geniculate (MGv) to AC projections. Recording electrodes were fabricated from borosilicate glass microcapillaries with a micropipette puller (Sutter). Before each whole-cell recording, the AC was identified by extracellular field responses to MGv stimulation.

#### Whole-cell current clamp recordings

Current-clamp recordings were obtained (Warner, PC-501A) to assess inhibitory postsynaptic potentials (IPSPs), excitatory postsynaptic potentials (EPSPs), and intrinsic firing rate in L2/3 AC pyramidal neurons. IPSPs were evoked via local biphasic stimulation of layer 4 (1 ms, 1-10 mV) in the presence of ionotropic glutamate receptor antagonists (20 μm DNQX; 50 μm AP-5). Antagonists were applied at least 8 min prior to IPSP recordings. Peak IPSP amplitudes of the short-latency hyperpolarization (putative GABAA component) and long-latency hyperpolarization (putative GABAB component) were measured from each response at a holding potential (Vhold) of −50 mV. EPSPs were electrically evoked from L2/3 pyramidal cells at a holding potential (Vhold) of −80 mV. EPSP waveforms were collected with increasing afferent stimulation levels (0.1 – 1 mA). Maximum amplitudes were determined from EPSP waveforms for each stimulation level to compute input-output functions. Action potential threshold was determined by delivering incremental current pulses (1500 ms; 10 pA steps; 0.2 Hz) until a spike was evoked. Frequency-current curves were collected by injecting depolarizing current steps (100-600 pA, 100 pA steps). Recordings were digitized at 10 kHz and analyzed offline using custom Igor-based macros (IGOR, WaveMetrics).

### Auditory brainstem response (ABR) measurements

ABRs were collected from all control and adolescent HL animals to determine whether the long duration of earplug insertion resulted in changes to the auditory periphery. ABR measurement procedures were identical to those used in previous studies in our laboratory (Rosen et al., 2012; Caras and Sanes, 2015; Yao and Sanes, 2018). ABRs were collected in animals following behavioral data collection and prior to electrophysiological experiments (see timeline in Figure 1A). Animals were anesthetized with intraperitoneal injections of ketamine (30 mg/kg; Bioniche Pharma) and pentobarbital (50 mg/kg; Sigma-Aldrich) and placed in a sound attenuation booth (Industral Acoustics). Recordings were made with subdermal needle electrodes inserted at the apex of the skull (positive electrode), the nape of the neck (inverting electrode; caudal to right pinna), and ground in hind leg. Sound-evoked ABR signals were preamplified (10,000x) and band-pass filtered (0.3-3 kHz) using a P55 model preamplifier (Grass Technologies). Signals were additionally amplified (32 dB gain; Brownlee 440 amplifier) before digitizing at a 24.4 kHz sampling rate (RZ6, TDT). Acoustic stimuli and data acquisition were controlled via custom Python scripts running on a PC (scripts provided by Brandon Warren and Edwin Rubel, University of Washington, Seattle). All stimuli were presented from a calibrated free-field speaker positioned 26 cm above the animal. Tone-evoked ABRs were collected by presenting tone pips (5 ms duration; 2 ms linear rise-fall times) with frequencies that spanned the range of the gerbil audiogram (0.5, 1, 2, 4, 6, 8, 16 kHz) (Ryan, 1976). Responses were averaged across 250-500 repetitions for each stimulus condition. Tone-ABR audiograms were generated by assessing the ABR threshold at each test frequency. We defined the ABR threshold to be the lowest sound level required to elicit a visually discernable ABR peak waveform. ABR waveforms exhibit stereotyped peaks that are representative of summed peripheral auditory nuclei electrical activity. Here, we focused on wave I of the ABR, which is thought to be generated by the auditory nerve (Jewett, 1970). We determined the peak of wave I amplitude, which is calculated as the absolute voltage difference between the peak and the following trough. Wave I amplitude was computed as a function of sound level to generate ABR input-output functions, which allowed us to assess whether earplugs induced subtle damage to auditory nerve synapses (Liberman and Kujawa, 2017).

#### Statistics

Statistical analyses and procedures were performed using JMP Pro 15.0 (SAS) or custom-written MATLAB scripts utilizing the Statistics and Curve Fitting Toolboxes. For normally distributed data (assessed using Shapiro-Wilk Goodness of Fit Test), values are given as mean ± SEM. Non-normally distributed data are given as median ± Quartile. Unless otherwise noted, all normally distributed data were assessed using parametric procedures (i.e., ANOVA) followed by appropriate *post hoc* controls for multiple comparisons (Tukey-Kramer HSD) when necessary.

## Abbreviations

(AC): auditory cortex
(HL): hearing loss
(AM): amplitude modulation
(IPSP): inhibitory postsynaptic potential
(EPSP): excitatory postsynaptic potential

## Acknowledgements

This work was supported by NIMH T32-MH019524 (KLA), NIDCD F32-DC018195-02 (KLA), and NIDCD R01-DC011284 (DHS). We would also like to acknowledge the Endocrine Technologies Core (ETSC) at the Oregon National Primate Research Center (ONPRC), which is supported by NIH Grant P51 OD011092 (ONPRC). Research reported in this publication through ETSC at the ONPRC was supported by the Office of the Director, National Institutes of Health of the National Institutes of Health under Award Number S10OD026701. The content is solely the responsibility of the authors and does not necessarily represent the official views of the National Institutes of Health. We also thank Lisa Ramirez for assistance in histological preparation and analyses. We are grateful to Melissa Caras and members of the Sanes lab for support and insightful feedback regarding analyses and interpretation of the data.

## Code availability

All behavior data were collected using custom MATLAB scripts developed by Dr. Daniel Stolzberg (https://github.com/dstolz/epsych). Psychometric fitting performed using Psignifit4 (https://github.com/wichmann-lab/psignfit/wiki). For *in vivo* physiology data: Semi-automatic spike sorting was performed using Kilosort (https://github.com/cortex-lab/KiloSort) and manually inspected and refined in Phy (https://github.com/cortex-lab/phy). For *in vitro* physiology data: All data were acquired using a custom-designed IGOR (version 4.08; WaveMetrics, Lake Oswego, OR) macro (Slice™). Software is available here: http://www.cns.nyu.edu/~sanes/slice_software/.

## Supplemental Figures

**Supplemental Figure 1.**
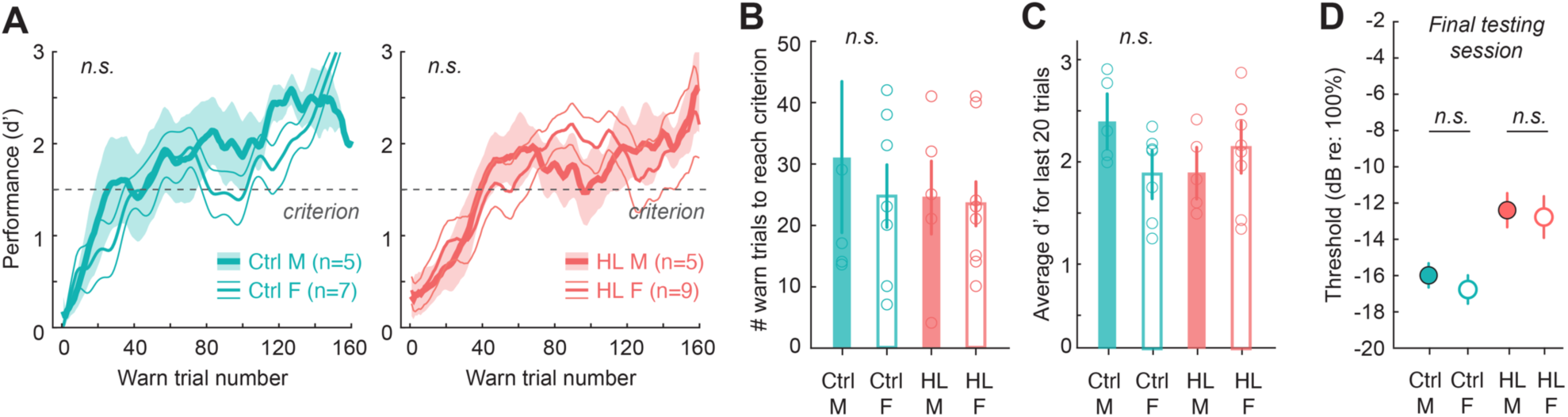
Transient hearing loss during adolescence did not alter procedural training or psychometric testing performance in a sex-specific manner. **(A)** Procedural training sessions (3-4 separate sessions) were combined separately for male (M) and female (F) control (Ctrl; left panel) and hearing loss-reared (HL; right panel) animals. Behavioral performance (d’) was computed as a function of warn trial number using a 5-trial sliding window. Data are depicted as the mean ± SEM. **(B)** Number of warn trials to reach performance criterion (d’≥; 1.5) for male (M) and female (F) control (Ctrl) and HL-reared animals. **(C)** Average d’ for the last 20 trials of procedural training for each group. **(D)** Control (Ctrl) and adolescent hearing loss (HL) males (M; closed circles) and females (F; open circles) exhibit comparable AM detection thresholds on the final testing session, with no significant differences between males or females within each group (controls: p=0.43; HL: p=0.97). Therefore, transient hearing loss during adolescence impairs amplitude modulation (AM) depth detection for both male and female adults.

**Supplementary Figure 2.**
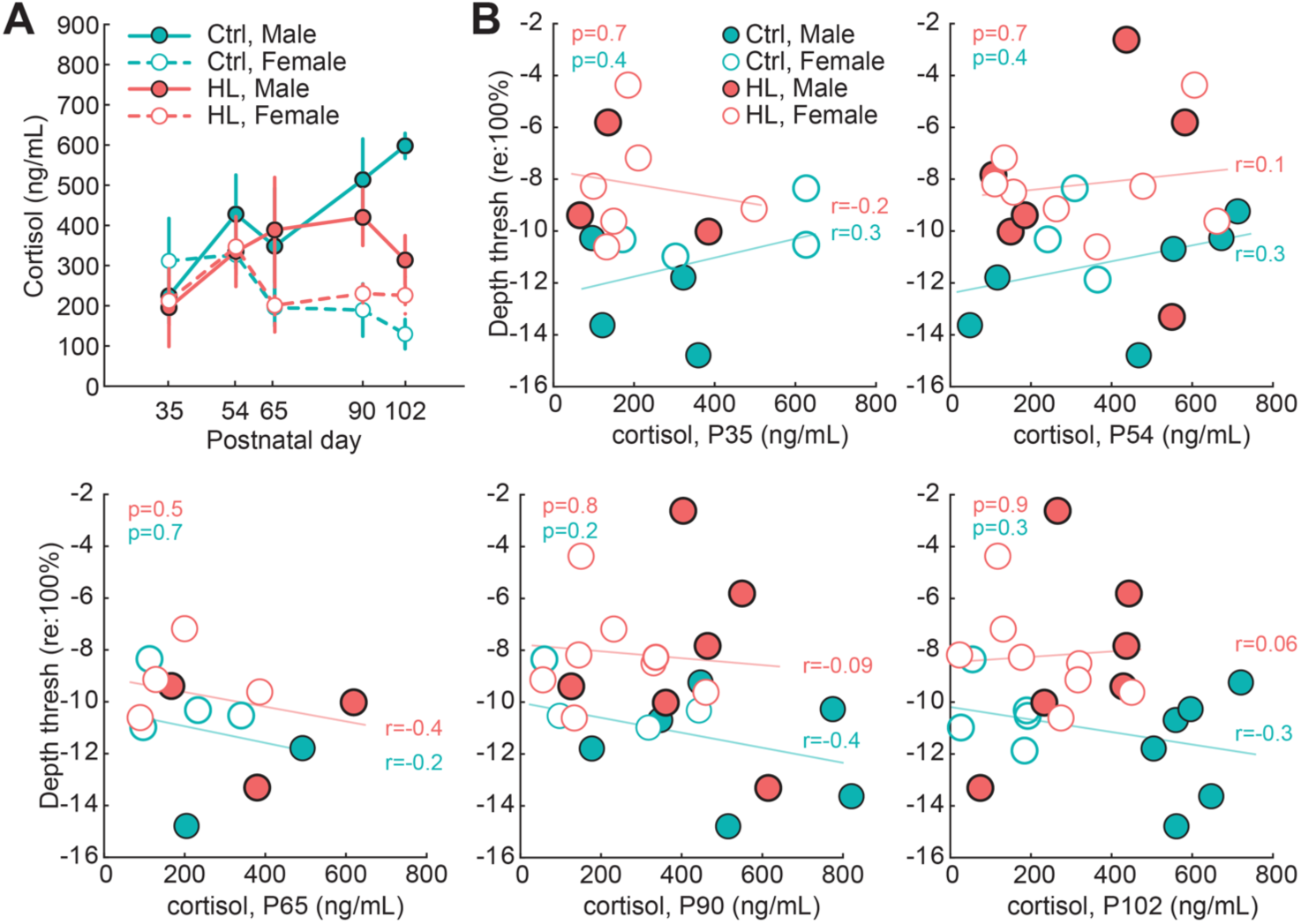
Poorer behavioral detection thresholds are not due to elevated stress during adolescence. **(A)** Serum cortisol levels across age for male (solid line) and female (dotted line) gerbils. **(B)** Behavioral depth thresholds for the first day of perceptual testing (∼P126; dB re: 100%) as a function of cortisol levels collected at P35, P54, P65, P90, and P102. Solid line indicates the linear fit for each group (Control, adolescent HL), along with the associated Pearson’s *r* correlation value. The p-values of each linear fit are listed in the top left corner of each plot. Elevated cortisol levels at any of the ages collected do not correlate with poorer detection thresholds at P126.

**Supplementary Figure 3.**
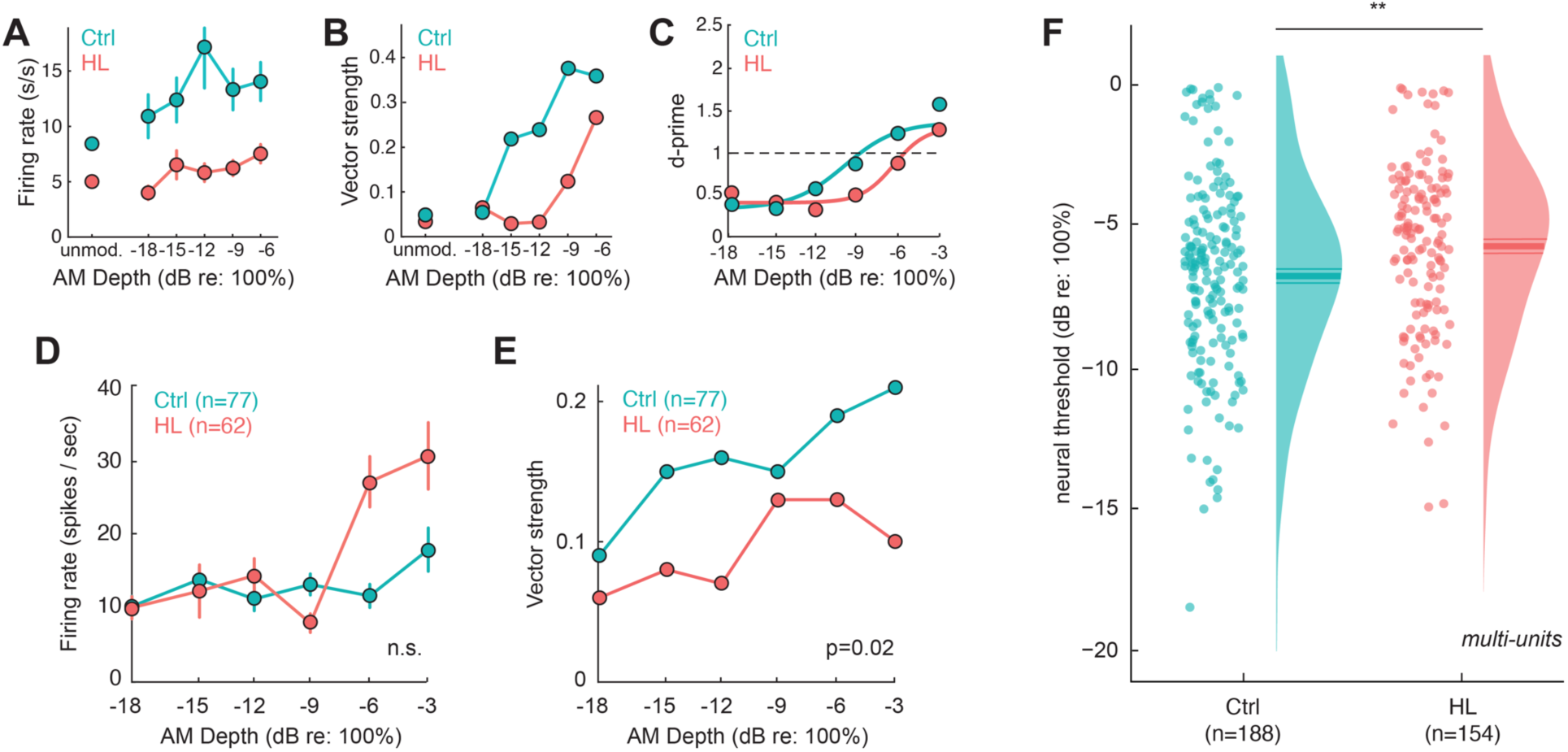
Basic response properties and detection thresholds for individual auditory cortex neurons. **(A-C)** Firing rate (s/s), vector strength (VS), and d’ values for example single units shown in Figure 4G. **(D-E)** Firing rate (s/s) and vector strength for a population of single units that met the criteria for AM sensitivity (same neurons from Figure 5). **(F)** Neural thresholds for multi units are plotted for control and HL animals. Individual thresholds are shown (circles), along with a half-violin plot indicating the probability density function. Horizontal lines indicate the mean±SEM. Individual multi units from HL animals exhibit poorer neural depth thresholds than control single units (p=0.003).

**Supplementary Figure 4.**
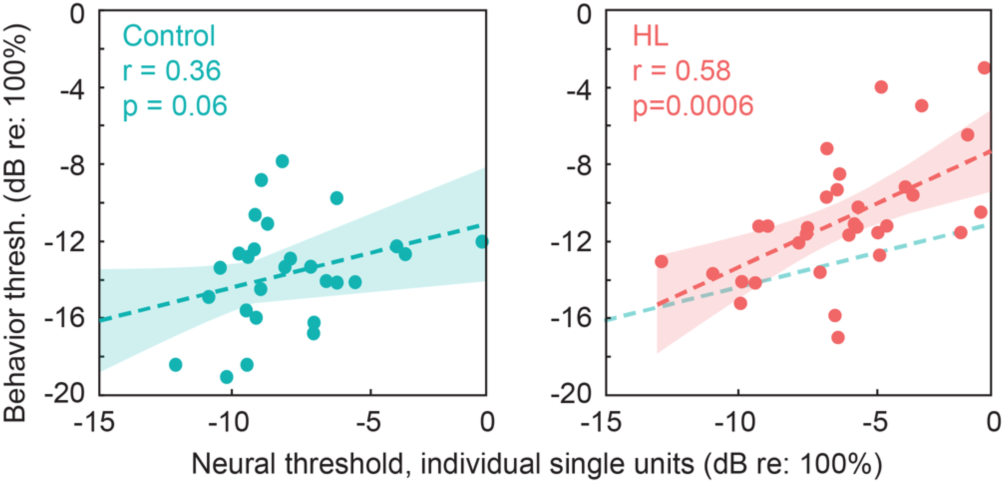
Neural sensitivity of individual cortical neurons correlate with perceptual performance. Behavioral threshold as a function of neural thresholds for individual single units that meet the criteria for AM sensitivity (see Methods). The neural thresholds are the average threshold for single units per session (i.e., 1 avg / session). Dotted lines indicate a fitted linear regression, with shaded areas indicating the (± 1 SD) of the prediction error. Pearson’s rand statistical significance of each fit are noted in the top left corner of each plot.

